# Functional traits predict outcomes of current and novel competition under warmer climate

**DOI:** 10.1101/2024.09.26.615168

**Authors:** Shengman Lyu, Jake M. Alexander

## Abstract

Functional traits offer a potential avenue to generalize and forecast the impacts of changing competition on plant communities, including changing outcomes of competition among species that currently interact (current competition) or that will interact in the future following range shifts (novel competition). However, it remains unclear how well traits explain variation in the outcomes of current and novel competition, as well as the underlying processes determining coexistence or competitive exclusion, under changing climate. Here, we conducted a field experiment in which pairs of high and low-elevation species interacted in three sites across an elevation gradient in the Swiss Alps. For each species pair, we quantified the population-level outcomes of competition (invasion growth rates), relative fitness differences and niche overlap and related these to 15 functional traits that were measured in each site. Most traits were significantly associated with invasion growth rates at the low elevation, where species had greater relative fitness differences, but these associations were much weaker towards higher elevations. This appears to be because traits, particularly those associated with light competition, captured species’ relative fitness differences at lower elevations, but not at the high elevation site. Greater relative fitness differences towards lower elevations suggest that climate warming may increase the likelihood of competitive exclusion of species that are poor competitors for light. In addition, novel competitors tended to show greater niche overlap than current competitors, leading to stronger overall competitive effects. But in general, trait differences predicted competitive outcomes of novel and current competitors similarly well, suggesting that traits can be used to predict interactions between species that do not yet interact. Our study reinforces the importance of considering changing interactions for predicting species responses to climate change and provides experimental evidence supporting the usefulness of functional trait differences in forecasting the impacts of future plant interactions under changing climate.

## Introduction

Climate change is reshaping global plant communities (Walther *et al*. 2002; Collins *et al*. 2022) directly by affecting species’ intrinsic population growth rates, and also indirectly by altering interactions between community members (Suttle *et al*. 2007; Gilman *et al*. 2010; Alexander *et al*. 2015). For instance, warming and altered rainfall can change the outcomes of competition between co-occurring species (Matías *et al*. 2018; Van Dyke *et al*. 2022). Over the longer term, species that previously did not co-occur will come into contact as species track climate change by shifting their ranges, giving rise to novel competitive interactions (Alexander *et al*. 2016; Descombes *et al*. 2020). A recent study shows that the expected upward migration of low-elevation species tracking climate warming into alpine communities could accelerate the local extinction of some alpine species through competitive displacement (Nomoto & Alexander 2021). Therefore, in order to accurately forecast species’ responses to climate change, it is critical to understand and predict how the impacts of both current and novel competitors vary under different climate scenarios (Gilman *et al*. 2010; Alexander *et al*. 2016). The plethora of possible interactions and the uncertainty in both future climate and the identities of novel competitors make it practically impossible to empirically measure the potential impacts of changing competition directly and generalize them across communities consisting of different sets of species. One possible solution to tackle this challenge is to link interaction outcomes to plant functional traits that characterize key aspects of the morphology, physiology, phenology and resource use of interacting species (Arnold 1983; Alexander *et al*. 2016; Díaz *et al*. 2016). However, our understanding of the impacts of competition under changing climate and our ability to predict them using functional traits remains limited for two main reasons.

First, past studies have tended to link interspecific trait differences to competition intensity measured on fitness components, most commonly on survival or biomass production (Violle *et al*. 2009; Kunstler *et al*. 2016; Lyu *et al*. 2017; Yang *et al*. 2022). However, effects on competition intensity offer limited insight into how trait differences influence the long-term population-level outcomes of competition because fitness components can respond to competition in varying or even opposing ways (Lyu & Alexander 2023). According to coexistence theory, the population-level outcomes of competition, that is, persistence or competitive exclusion, can be captured by species’ abilities to grow from low density (invasion growth rates) while their competitors are at equilibrium densities (Chesson 2000; Grainger *et al*. 2019). Additionally, coexistence theory emphasizes two fundamental processes determining the outcomes of competition (Chesson 2000; Spaak & De Laender 2020). One is niche differentiation (i.e., reduction in niche overlap), arising from, for example, resource partitioning (Tilman 1982), that can promote stable coexistence. The other is relative fitness differences, which reflect the extent of the asymmetry in species’ competitive abilities that drive competitive exclusion. A focal species might be able to persist with a competitor because it possesses trait values that confer upon it a competitive advantage and/or that reduce its niche overlap with the competitor (Carroll *et al*. 2011; Ellner *et al*. 2016; Grainger *et al*. 2019). In sum, establishing associations between trait differences and invasion growth rates provide a theoretically justified test for the ability of functional traits to predict the outcomes of competition. In addition, examining associations of traits with relative fitness differences and niche overlap can provide deeper insights into the processes that regulate the impacts of competition and thereby help generalize the predictive ability of functional traits (Adler *et al*. 2013).

A second hindrance to using functional traits to understand and predict competition under changing climate is that most recent studies have linked functional traits to competition in a single environment and between currently co-occurring species (e.g. Kunstler *et al*. 2012; Kraft *et al*. 2015). However, the relationships between traits and species coexistence may vary with climate and between current and novel competitors. First, the relevant traits affecting interaction outcomes might change if the main limiting resources shift as climates change (Copeland & Harrison 2017; Borges *et al*. 2019; Perez-Ramos *et al*. 2019; Li *et al*. 2022). For instance, traits associated with access to light, such as plant height and leaf economics traits (Adams *et al*. 2007), may have great impacts on competition under light-limited conditions, such as in warm climate where productivity is usually high, but have little impact under conditions where light is not limiting, such as in cold climates where productivity is limited by low temperatures (Walker *et al*. 2006; Bjorkman *et al*. 2018). Even if the primary limiting resources are similar, species likely display greater trait differences under productive (e.g. warm) than limiting (e.g. cold) climate (Wei *et al*. ; Sherry *et al*. 2007) and thus are likely to be more differentiated along a competitive ability hierarchy or within niche space (Matías *et al*. 2018), leading to closer relationships between trait differences and coexistence.

In addition to environmental variation, trait-competition relationships might also differ between novel and current competitors for at least two reasons. First, novel competitors that originate from distinct ecoregions and divergent lineages are expected to possess greater trait differences than co-occurring species that have undergone shared abiotic and biotic filtering (van Kleunen *et al*. 2009; Alexander *et al*. 2015; Mathakutha *et al*. 2019; Zhang & van Kleunen 2019). In this case, amplified trait differences, particularly if they are associated with relative fitness differences, might increase the strength of associations between traits and coexistence for novel versus current competitors. Secondly, the long-term history of co-occurrence between current competitors may have led to character displacement that favours their stable coexistence (Zuppinger-Dingley *et al*. 2014; Germain *et al*. 2018b; Sakarchi & Germain 2023). Recent theoretical studies suggest that character displacement can facilitate the stable coexistence of competing species either via the divergence of traits associated with niche use (reducing niche overlap), or via the convergence of traits associated with species’ competitive abilities (reducing relative fitness differences) (Germain *et al*. 2018b). Such adjustments might weaken associations between traits and competitive outcomes in the case of current competitors. For instance, plant height might have strong impacts on the interactions between novel competitors due to competition for light, but have weaker impacts on interactions between current competitors if they have evolved ways to avoid height-mediated light competition, such as through divergence in phenology (Blackford *et al*. 2020).

In this study, we conducted a field experiment in which pairs of low and high-elevation species (lowland and highland species) interacted in three sites across an elevation gradient in the Swiss Alps, where climate change and associated elevational range shifts have been documented (Rumpf *et al*. 2018; Vitasse *et al*. 2021). Elevation gradients are useful systems to study the impacts of climate change on plant communities, where a series of abiotic (e.g. temperature, soil moisture and their covariation) and biotic (e.g. competitors) factors systematically vary across space in a way that is analogous to expected climate change over time (Elmendorf *et al*. 2015). We selected seven lowland and seven highland species that are common in their respective communities but have little range overlap and thus limited co-occurrence history. Therefore, interactions between lowland-lowland or highland-highland species reflect current interactions (n = 56 pairs), while interactions between lowland-highland species reflect novel interactions (n = 56 pairs). We parameterized integral projection models (IPMs) to predict population-level outcomes of competition (i.e. invasion growth rates)(Grainger *et al*. 2019) and quantified relative fitness differences and niche overlap for competitor pairs at each site. We measured 15 traits encompassing morphology, physiology, phenology, and resource use for each species at each site (Table 1). We then related interspecific trait differences to the invasion growth rates and the two coexistence determinants and examined how these relationships varied across the elevation gradient. With these analyses, we aim to increase our understanding of the impacts of climate change-induced alterations to competition on plant communities and provide a rigorous experimental test of the usefulness of traits in predicting these impacts. Specifically, we ask: (1) Are interspecific trait differences associated with invasion growth rates and the two coexistence determinants? (2) How do the relationships between trait differences and coexistence vary across the elevation gradient? (3) How do the relationships between trait differences and coexistence differ between current vs novel competitors?

**Table 1.**
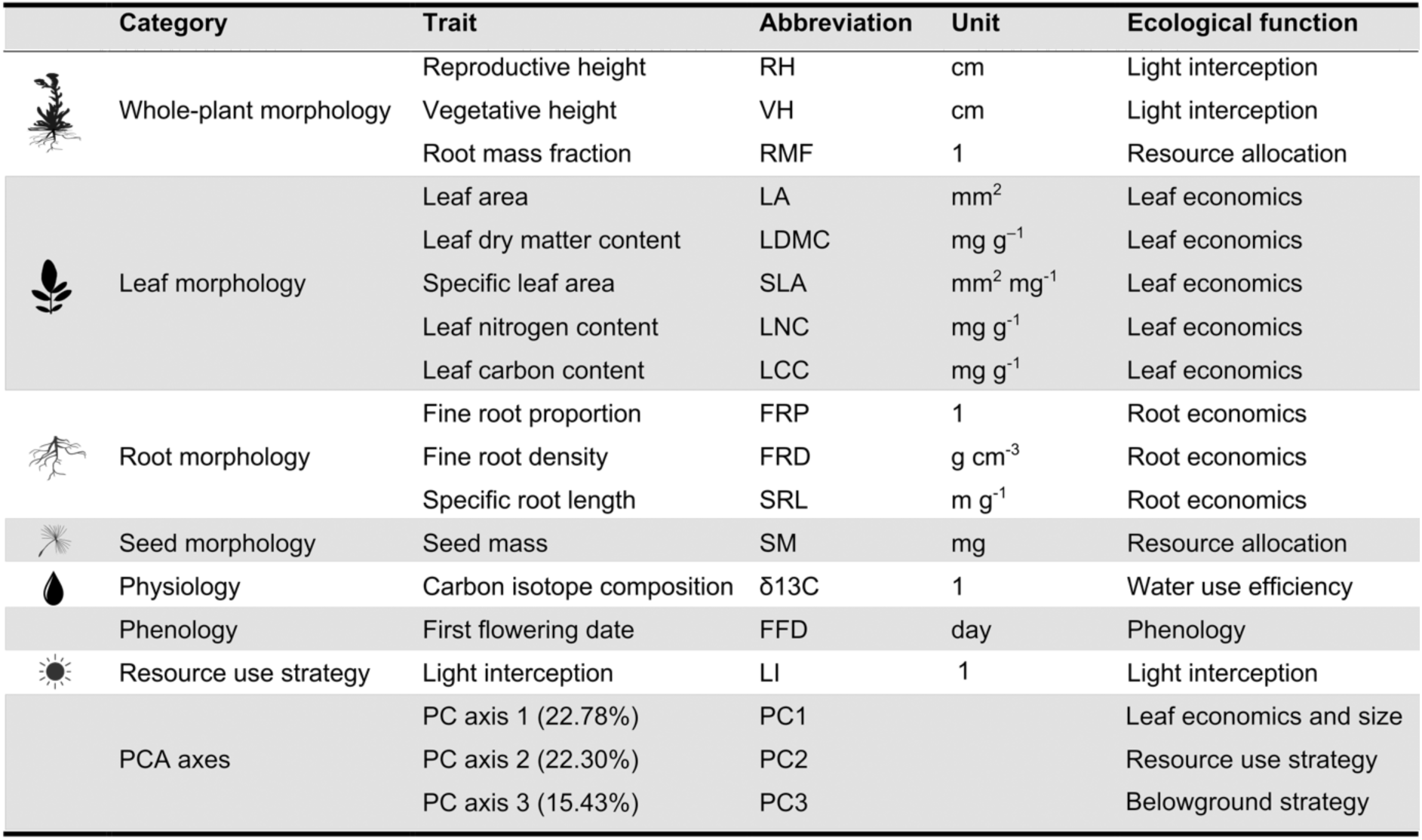
Species’ traits included in this study, with their abbreviations, units and associated ecological functions. The values by PCA axes indicate the percentage of the total variation for which each axis accounts.

## Materials and methods

### Study sites and species

We conducted a field experiment across an elevation gradient in the Swiss Alps (46°30′N; 7°10ʹE), including three sites located at 900, 1400 and 1900 m above sea level (hereafter, the low, middle and high sites). The three sites represent a climate gradient with long-term (1981 to 2015) mean annual temperatures ranging from 2.5, 5.9 to 9.7 °C at the high, middle and low sites, respectively (Scherrer *et al*. 2020). The three sites were located on south-facing shallow slopes (less than five degrees) in summer pastures (no woody plants) with grazing excluded throughout the experiment, allowing us to focus on the effects of climate on species interactions. We selected 14 common perennial species originating from the low and high-elevation communities (hereafter, the lowland and highland species; Table S1). Lowland species shared upper range limits below 1500 m, defined as the 90% quantile of their elevation range, except for *Plantago lanceolata* with an upper range limit of 1657 m; highland species shared lower range limits above 1500 m, defined as the 10% quantile of their elevation range based on a vegetation dataset collected within the study area (Randin *et al*. 2009). Therefore, the lowland and highland species had limited co-occurrence history. To avoid biasing our species selection towards greater trait variation between lowland and highland species than within each group (e.g., only tall lowland species and short highland species), we selected lowland and highland species to display a wide range of traits within each group based on plant height, specific leaf area and seed mass derived from the LEDA dataset (Kleyer *et al*. 2008; Fig. S3).

### Field experiment

We designed a field experiment to quantify pairwise outcomes of competition across the elevation gradient and between species that currently co-occur (i.e. current interactions, lowland-lowland and highland-highland species pairs) and do not co-occur (i.e., novel interactions, lowland-highland species pairs). Within each site, each species interacted with four current and four novel competitors, giving rise to 112 (14 x 8) interspecific pairs with 56 current and 56 novel pairings. We selected these interspecific pairs to evenly sample the functional trait difference of all possible pairwise current and novel combinations based on their plant height, specific leaf area and seed mass. In addition, each species also interacted with itself (*n* = 14 intraspecific pairs) and grew in the absence of neighbours (*n* = 14 non-competition) at each site. We had nine focal individuals for each species pair, giving rise to 3780 individuals in total in the full design [(56 current pairs + 56 novel pairs + 14 intraspecific pairs + 14 non-competition) x 9 individuals x 3 sites].

In spring 2017, we established 18 plots (1 x 1.6 m and 0.2 m deep) within each site and lined them with wire mesh to exclude rodents (except for the high site) and with weed-suppressing fabric on the sides to prevent plants from growing in from outside. We filled all plots in the three sites with soil originating from a location near the low site, allowing us to isolate the effects of climate on species interactions from possible effects of soil. We then sowed each species into a plot with a density of 9 g m^-1^ viable seeds as background competitors (*n* = 14 competition plots) and left the other plots as bare soil (*n* = 4 non-competition plots). We periodically weeded the plots throughout the experiment to maintain background monocultures.

We raised seedlings in a greenhouse and transplanted them into the field sites as focal plants. We used standard compost soil and let seedlings grow in the greenhouse for six weeks and acclimate in a common garden for a week before transplanting. In autumn 2017, we transplanted focal seedlings into established competition plots at 14 cm spacing and non-competition plots at 25 cm spacing. Focal seedlings that died within two weeks after transplanting were replaced (ca. 5%). We had to resow species that failed to establish, and they received focal plants either in spring 2018 (*Poa trivialis* and *Poa alpina* at the low site and *Bromus erectus* at the middle site) or autumn 2018 (*Aster alpinus*, *P. trivialis* and *P. alpina* at the middle site and *Sesleria caerulea* at the low and high sites). We replaced dead focal individuals in the spring and autumn of 2018 and 2019 (ca. 10% each time). We included species that failed to establish as competitors only as focal species for calculating population growth (*Daucus carota* at all sites and *S. caerulea* at the middle site).

### Vital rate data collection

We monitored the vital rates of all focal plants between 2017 and 2020. Survival was recorded twice a year, at the beginning and end of each growing season. Towards the end of the growing season when all species were fully grown and flowering (between July and September depending on the site), we measured size-related morphological traits (including the number and length of flowering stems, the number and length of leaves or ramets, depending on the species), recorded flowering and counted the number and measured the size of fruits or flowers. To estimate plant size, we fitted linear models of dry aboveground biomass as predicted by the same set of morphological traits, using plants collected from the background monocultures. To estimate fecundity, we fitted linear models of the number of seeds as predicted by the size of fruits or flowers, using intact fruits collected from background monocultures at early fruiting stages. We conducted a separate experiment to estimate seed germination and seedling establishment in the absence of neighbours. We also estimated competition-dependent seedling establishment as the survival probability of focal seedlings within their first growing season after transplanting. See (Lyu & Alexander 2022) for further details on demographic data collection.

### Population modelling

We used integral projection models (IPM) to model population growth (Ellner *et al*. 2016). An IPM incorporates vital rate transitions from census *t* to *t* + 1 into population growth, as denoted by the integral:

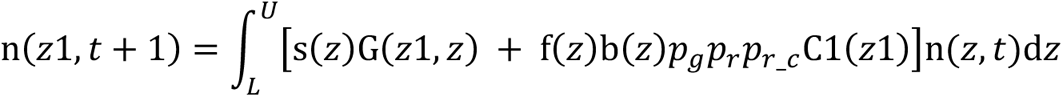

where, *z* represents plant size (i.e., dry aboveground biomass on a natural logarithm scale) and was used as a continuous state variable in the IPMs. n(*z*, t) represents the population size distribution at census t. *L* and *U* represent the lower and upper size bounds covering all possible sizes within the population. Vital rate functions describe transitions from census *t* to *t* + 1. These transitions were either dependent on size, z, including probability of survival [s(*z*)], growth [G(*z*1,*z*)], probability of flowering [f(*z*)], fecundity [b(*z*)], and offspring size distribution [C1(*z*1)], or independent of size, including seed germination rate (p_g_), competition-free establishment probability (p_r_) and competition-dependent establishment probability (p_r_c_). Further details on model structure and parameters can be found in (Lyu & Alexander 2022).

We combined all three censuses, that is, 2017-2018, 2018-2019, and 2019-2020, to parameterise the IPMs and estimate deterministic population growth rates (λ). To examine the effects of plant size, competitor species and site on each vital rate of each species, we used a model selection approach to compare all the nested models of the full models including all three factors and their interactions. The full models of size-independent vital rates, that is, seed germination and seedling establishment, included only competitor species, site and their interactions. We compared all candidate models using the Akaike information criterion corrected for small samples (AICc). We used simplified models in cases where the lowest-AICc model appeared to be overfitting based on visual checks. We then obtained vital rate parameters from the best-fit vital rate.

We calculated deterministic population growth rates (λ) as the dominant eigenvalue of the parameterised IPMs (Ellner *et al*. 2016). We estimated intrinsic population growth rates (λ_intrinsic_) based on IPMs fitted using plants growing in the absence of neighbours and the low-density population growth rates (λ_invasion_) based on IPMs fitted using plants invading the established monocultures of competitors. We had to exclude *Arnica montana* at all sites and *Trifolium badium* at the low site for the calculation of λ due to data scarcity caused by high mortality rates.

### Outcomes of competition, relative fitness differences, and niche overlap

We used λ_invasion_ as a measure of species’ persistence under competition (i.e. the outcomes of competition). An λ_invasion_ greater than one indicates that the population is predicted to persist in the presence of neighbours, with greater values indicating a greater ability to persist. An λ_invasion_ less than one indicates the species is predicted to be competitively eliminated by its competitor.

We also quantified niche overlap (NO) and relative fitness differences (RFD; i.e. competitive ability differences) based on the estimates of population growth rates in the absence and presence of neighbours (Carroll *et al*. 2011; Narwani *et al*. 2013). For a focal species *i* competing against species *j*, we first calculated its sensitivity as:

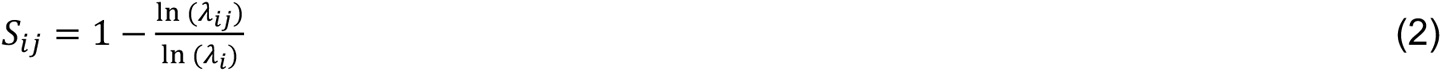

where λ*_ij_* is the invasion growth rate of species *i* in the presence of its competitor *j*, which is at its equilibrium density, and λ*_i_* is its intrinsic growth rate. Note that this equation assumes the monocultures of competitor *j* is at its equilibrium density (see a test for this assumption in Lyu & Alexander 2022). Sensitivity is positive for competitive interactions, and greater sensitivities indicate stronger competition; sensitivity is negative for facilitative interactions.

Modern coexistence theory predicts the outcomes of competition are determined by the relative magnitude of niche overlap and relative fitness differences. Niche overlap hinders species coexistence because it increases the intensity of interspecific competition (Chesson 2000; Godoy & Levine 2014). Therefore, a pair of species that experience more intense interspecific competition (i.e., greater mean sensitivities) have greater niche overlap, as captured by the geometric mean of sensitivities:

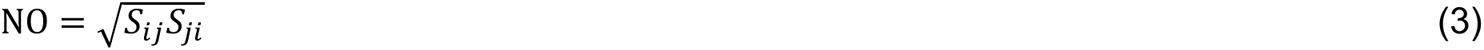

Relative fitness differences quantify the degree of asymmetry in species’ competitive abilities and drive competitive exclusion (Chesson 2000; Godoy & Levine 2014). Therefore, a pair of species that experience different levels of interspecific competition (i.e., greater differences in sensitivities) tend to have great relative fitness differences, as captured by the geometric standard deviation of sensitivities:

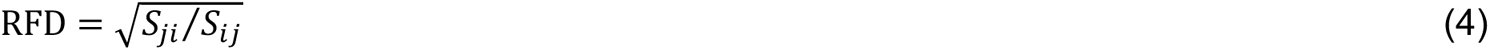

There are three possible outcomes of competition determined by the relative balance of niche overlap and relative fitness differences (Chesson 2000). Competitive exclusion occurs when two species have high niche overlap and large fitness differences, and stable coexistence occurs when niche overlap is less than 1 (intraspecific competition > interspecific competition) and small enough to overcome the impacts of the fitness difference driving competitive exclusion. Otherwise, when species have high niche overlap and small relative fitness differences priority effects occur, where the species initially established within a community excludes the other. Equations 3 and 4 show that this approach is incompatible with facilitative interactions whose sensitivities are negative; thus, we included only pairs that experienced competitive interactions for the analyses (92% of 284 pairs). The results were very similar when we included facilitative interactions in the analyses by scaling their sensitivities to 0.1 (indicating weak competition).

### Trait measurements

We measured 15 traits on plant morphology, physiology, phenology, and resource use within the field experiment. These traits represented various hypothesized ecological functions (Table 1), which were measured following standard protocols (Perez-Harguindeguy *et al*. 2013). We measured plant reproductive and vegetative heights on randomly selected individuals from the background monocultures when plants were fully grown in 2018 and 2020 (*n* = 10 to 20 individuals per species per site).

To measure leaf traits, we collected leaf samples at peak growing season between 2018 and 2020 (*n* = 30 to 50 samples per site per species), kept them moist and cool, and measured fresh mass and scanned leaf area with a scanner (CANON CanoScan LiDE 400) within 24 hours of collection. These leaf samples were dried at 70°C for 72 hours and weighed for dry mass. We used leaf samples collected in 2020 (*n* = 6 samples per species per site) to measure leaf nitrogen and carbon content and carbon isotope composition with an isotope ratio mass spectrometer (IRMS, DeltaplusXP, Finnigan MAT, Bremen, Germany).

To measure seed mass, we collected mature seeds from background plants at the end of the growing season in 2019 and 2020, which were then air-dried and cleaned. We mixed seeds from multiple individuals within each site and counted 10 to 100 seeds depending on the seed size and weighed for air-dried mass. Seed mass was calculated as the total weight divided by the number of seeds (average of *n* = 4 to 6 samples per species per site).

To measure root traits, we harvested six individuals per species per site within background monocultures at the end of the growing season in 2020. The aboveground parts were separated and oven-dried for dry shoot biomass. The entire roots were carefully washed over a 500 nm sieve under running water and stored in 70% ethanol before further measurements. We scanned the whole root floating in water with a scanner (Epson Expression 10000XL) and analysed root diameter and length with WinRhizo software (basic version 2007a). We followed the same approach to measure the length and volume of fine roots (diameters < 2 mm) of each sample, which were then dried at 70°C for 72 hours and weighed for dry mass.

To monitor flowering phenology, we estimated the proportion of flowering plants (i.e., plants with flowering buds or open flowers) within each plot weekly between April and September in 2019. We fitted generalized additive models (GAM) with a binomial distribution of the proportion of flowering individuals as explained by the Julian date of the year. We then predicted the first flowering date using the fitted GAMs (i.e., the Julian date when the proportion of flowering plants shifted from zero to positive). We used the GAM-derived first flowering date rather than the actual date when flowers were first observed because we could miss the start of the flowering of one species with the earliest flowering phenology (*Sesleria caerulea*), and also to reduce the effect of outliers (e.g., by plants that flowered extremely early).

To characterise species’ ability to intercept light, we measured light interception of background monocultures as the degree to which each species drew down ambient light. At peak growing season in 2019, we measured light intensity above (ca. 1 m above the ground surface) and below the canopy at ten haphazardly chosen locations within each plot (but avoiding the influence of focal plants) with a photometer (SDEC, France; www.sdec-france.com). The light intercept was calculated as the percentage of ambient light getting through the canopy, with the smaller value indicating a greater ability to intercept light.

### Statistical analyses

Prior to analyses, we averaged each trait to obtain a mean value per species and site (Fig. S1). The site-level mean of traits was log or square root transformed to improve the symmetry of distributions and standardized to have a mean of zero and a standard deviation of one across all species and sites to facilitate comparison between traits.

To test the effects of individual traits on the outcomes of competition (i.e. invasion growth rates), we fitted a linear mixed-effects model for each trait separately with λ_invasion_ (on a natural logarithm scale) as the response variables, hierarchical trait differences (trait_focal_ – trait_competitor_), site (categorical variable, low, middle or high), the origin of competitor (current or novel), and all two-way and three-way interactions as explanatory variables, and the identity of focal and competitor species as random effects. The three-way interactions were then removed since they were insignificant in any models. We tested the significance of fixed effects retained in the simplified models using type-II *F*-tests. We then extracted the slopes of hierarchical trait difference to determine whether traits had significant effects on λ_invasion_ at a certain site or for a certain of competitor origin. We calculated the 95% profile confidence intervals of the slopes using the *confint.merMod* function in the R package lme4 and determined the significant effects of a trait if the 95% CI of its slope did not include zero. Similarly, we also tested the impacts of interspecific trait differences on relative fitness differences and niche overlap. We used hierarchical trait differences (trait_focal_ – trait_competitor_) for relative fitness differences, assuming competitive dominance to be directional, and absolute trait differences (|trait_focal_ – trait_competitor_|) for niche overlap, assuming it to be non-directional (Kraft *et al*. 2015; Pérez-Ramos *et al*. 2019).

Because multiple biological processes can operate simultaneously to determine the outcomes of competition, we also tested the effects of interspecific differences in multiple traits on coexistence. First, we performed a principal component analysis (PCA) with all traits and species across the three study sites. We then extracted the PCA scores for each species in each site along the first three PCA axes. We calculated interspecific differences using PCA scores to measure multivariate trait differences between species. We fitted linear mixed-effects models for invasion growth rate, relative fitness differences and niche overlap separately for each PCA axis, similar to the models with individual traits. Second, we also performed multivariate analyses to test whether multiple traits in specific combinations that were not captured by the PCA axes affect coexistence (see detailed description in the Supplementary methods). We log-transformed the response variables and visually verified the model assumptions (linearity, homogeneity of variance and Gaussian error distribution) in all models. All population modelling and statistical analyses were performed using R version 4.0.3 (R Core Team 2020).

## Results

### How do outcomes and competition, relative fitness differences and niche overlap vary across the elevation gradient and between current and novel competitors?

Among all 93 species pairs for which the outcomes of competition were predicted across the three study sites, approximately two thirds of them (n = 61) were predicted to be able to stably coexist (Fig. 1). The frequency of coexisting pairs tended to be lower at low elevation, with 55%, 82%, and 61% at the low, middle and high sites, respectively (Fig. 1 a-c). The greater frequency of competitive exclusion at low elevation appeared to be mainly driven by amplified relative fitness differences (F_2, 93_ = 318.785, P < 0.0001; Fig. 1 d), while niche overlap did not change significantly with elevation (F_2, 93_ = 2.007, P = 0.367;Fig. 1 e). In addition, current competitors in general had greater abilities to coexist than novel competitors, with 74% of current pairs and only 56% of pairs predicted to be able to coexist (Fig. 1 a-c). This was mainly because current pairs had smaller niche overlap than novel pairs (F_1, 93_ = 4.985, P = 0.025; Fig. 1 e), while current and novel pairs had similar magnitudes of relative fitness differences (F_1, 93_ = 0.29, P = 0.59; Fig. 1 d).

**Figure 1.**
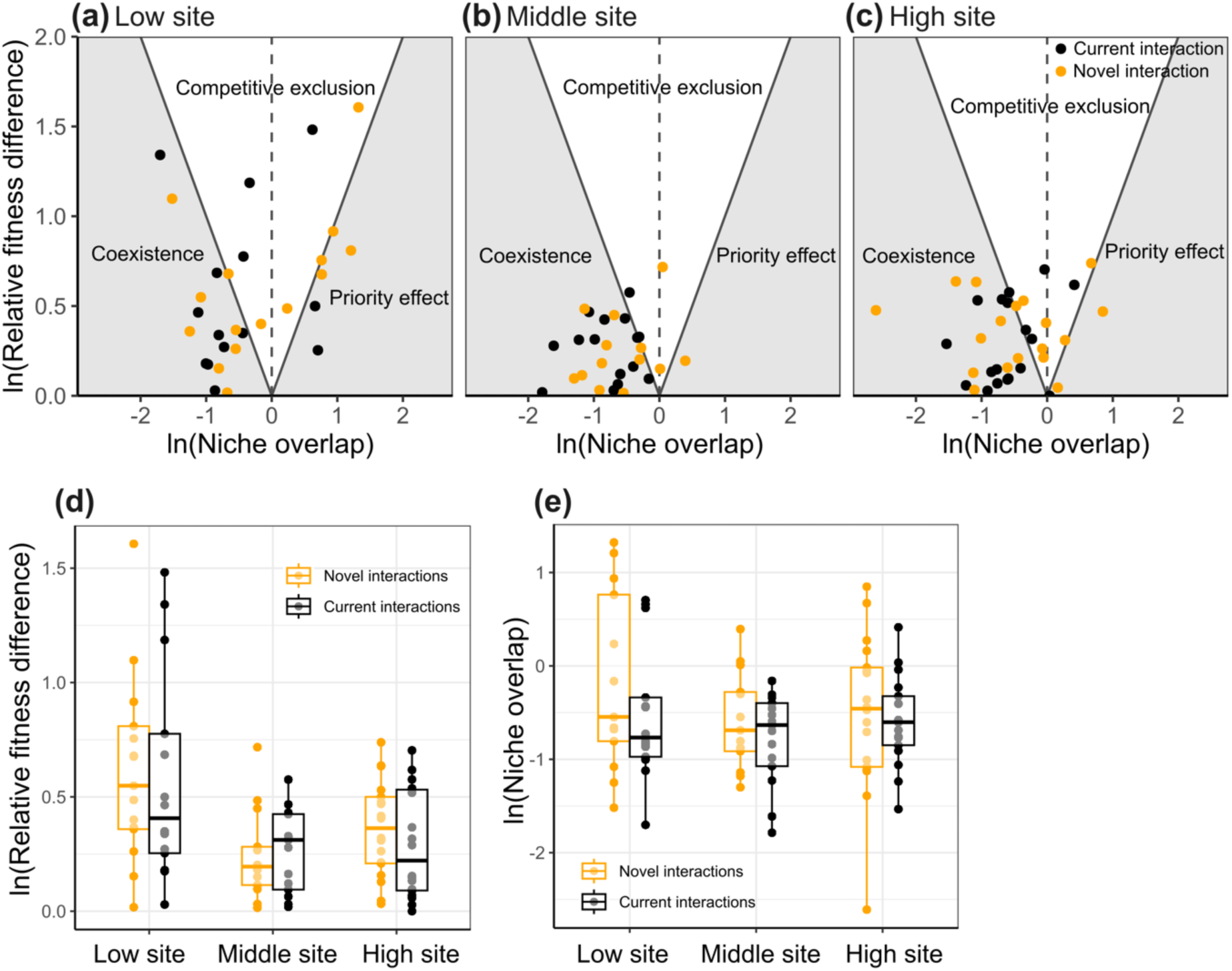
Coexistence in the three study sites across an elevation (low site, 890 m; middle site, 1400 m; high site, 1900 m). Panels a-c are the outcome of competition for current (black) and novel (orange) interactions at the low (a), middle (b) and high (c) sites. Each dot represents a species pair. Each panel shows three possible outcomes: stable coexistence (shaded area when niche overlap is negative), priority effects (shaded area when niche overlap is positive), and competitive exclusion (unshaded area). Panels d-e show relative fitness differences (d) and niche overlap (e) for current (black) and novel (orange) interactions across the three study sites. Boxplots represent the median, first, and third quartiles summarized across species pairs within each site, while the upper and lower whiskers indicate 1.5 times the first and third quartiles, respectively.

### Are interspecific trait differences associated with invasion growth rates and the two coexistence determinants?

We found that interspecific trait differences were frequently related to the outcomes of competition. Among the 15 traits we measured, 12 traits were significantly related to invasion growth rates in at least one site (Fig. 2 and Fig. S7 in the Supplementary Information). These showed that relative to competitors focal plants with greater plant height, larger and thicker leaves, lower leaf nitrogen and carbon concentration, more coarse roots, denser fine roots, heavier seeds, or greater abilities to intercept light had a greater ability to persist under competition compared with other species (Fig. S7). Of the two determinants of the outcome of competition, functional traits were more frequently related to relative fitness differences than niche overlap (Fig. 3). Specifically, 11 traits were significantly related to relative fitness differences across the three study sites (Fig. 3 a&b). Those showed that species had greater competitive ability when they were taller or had greater biomass allocation to shoots, larger leaves or leaf dry matter content, lower density of fine roots, less leaf carbon, heavier seeds, higher water-use efficiency, or later flowering phenology. In contrast, only seven traits across the three study sties were significantly related to niche overlap (Fig. 3 c&d); species had greater niche overlap when they had more similar leaf area, leaf dry matter content and nitrogen concentration, specific root length, seed mass, flowering time, and light interception ability compared to species with more distinct trait values.

**Figure 2.**
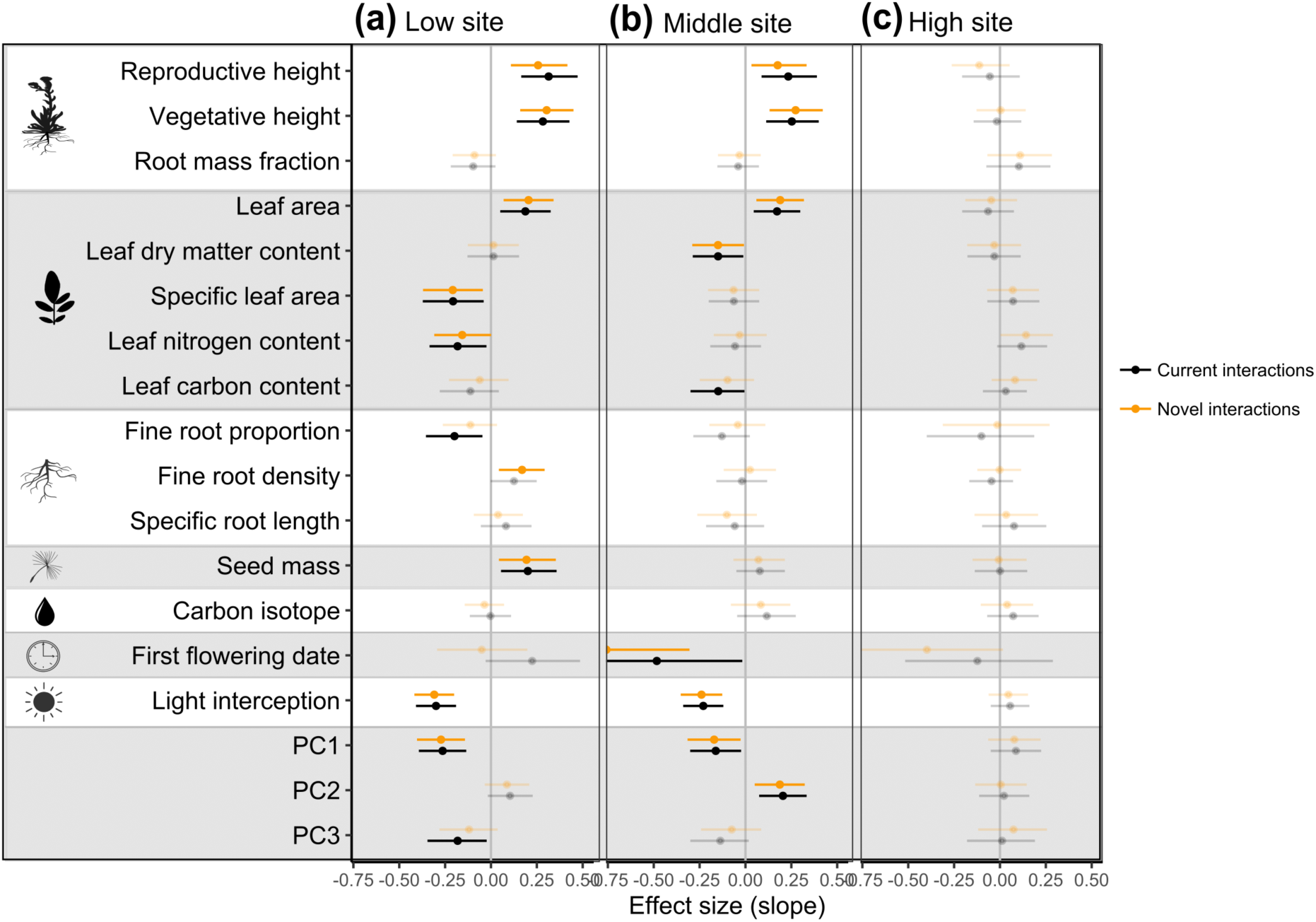
Effect sizes of each trait on invasion population growth rates in the three sites across the elevation gradient (a, low site, 890 m; b, middle site, 1400 m; c, high site, 1900 m) for current (black) and novel (orange) competitors. Effect sizes are slopes of the relationships between interspecific trait difference (i.e., trait_focal_ – trait_competitor_) and invasion population growth rates (i.e., λ_invasion_; Fig. S8). Positive effect sizes suggest that, relative to competitors, focal species with greater trait values have greater ability to persist under competition (Fig. S8). Points and error bars represent the estimated effect sizes and their 95% CIs. Effect sizes are determined as significant if the 95% CI do not include zero (opaque dots and error bars).

**Figure 3.**
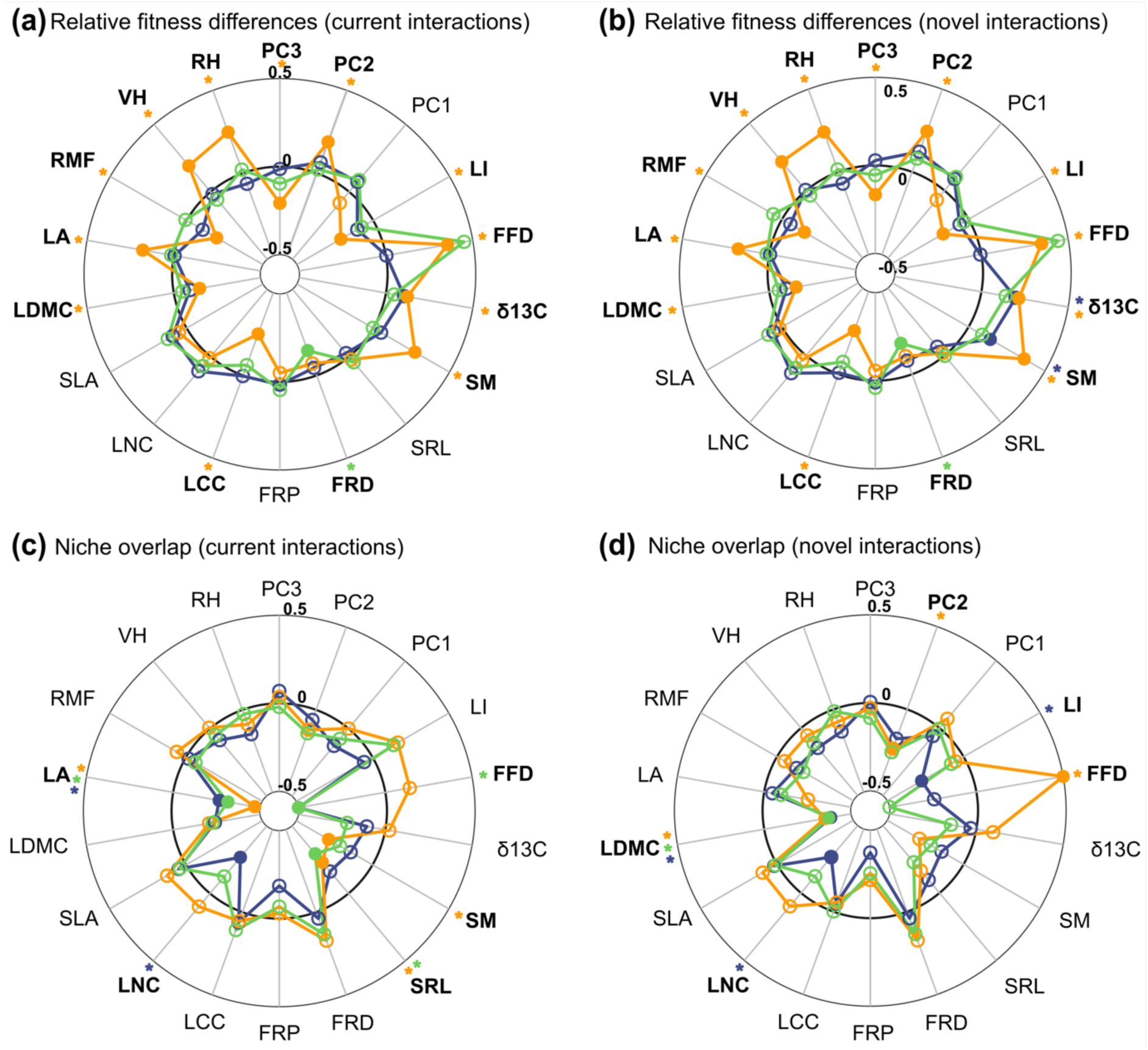
Effect sizes of each trait on relative fitness differences (a and b) and niche overlap (c and d) between current (a and c) and novel (b and d) competitors in the three sites across the elevation gradient (orange, low site, 890 m; green, middle site, 1400 m; blue, high site, 1900 m). The inner, middle, and outer cycles indicate the effect sizes of -0.5, 0 and 0,5, respectively. For relative fitness differences in A and B, positive effect sizes suggest that, relative to competitors, focal species with greater trait values are competitively dominant. For niche overlap, negative effect sizes suggest that competitors with more different trait values have smaller niche overlap. Effect sizes that are significant (the 95% CI does not include zero) are indicated with filled dots and asterisks on the outer cycle, while insignificant effects sizes are indicated with unfilled dots.

These effects on competition of individual traits were mirrored in the multidimensional trait variation captured by the PCA. PC1 (23% of total variance), which mainly reflected the leaf economics spectrum (Wright *et al*. 2004) and variation in aboveground size (Fig. S3), was negatively related to invasion growth rates. This indicated that species possessing more conservative strategies, such as lower specific leaf area and leaf nitrogen concentration, heavier seeds, and larger size than their competitors, were better able to persist under competition (Fig. 2). Traits associated with acquisitive resource use aboveground, such as high light interception and long-term water use efficiency, loaded heavily on PC2 (22% of variance). The positive effect of PC2 in the middle site indicated that species that were more efficient at resource acquisition and use were better able to persist under competition (Fig. 2). Additionally, a negative effect of PC3 (15% of variance), which mainly captured belowground strategies, in the low site showed that species with more conservative strategies (such as higher root density and root mass fraction but lower specific root length) relative to its competitor were better able to persist under competition (Fig. 2). In terms of the two processes determining competitive outcomes, PC2 (capturing aboveground resource use strategy) and PC3 (capturing belowground strategies) were significantly related to relative fitness differences (Fig. 3 a&b), while only PC2 was significantly related to niche overlap (Fig. 3 d).

Among the 12 individual traits that significantly related to invasion growth rates across the three study sites, five were significantly related to both relative fitness differences and niche overlap (LA, LDMC, SM, FFD and LI; Fig. 3), while a further four traits were only significantly related to relative fitness differences and an additional one trait related to niche overlap (Fig. 3). The other two traits, SLA and FRP, were not significantly related to relative fitness difference or niche overlap (Fig. 3), despite their significant relationships with invasion growth rates (Fig. 2).

### How do relationships between trait differences and coexistence vary across the elevation gradient?

The relationships between interspecific trait differences and invasion growth rates varied greatly across the elevation gradient. Among the 12 individual traits significantly related to invasion growth rates, 11 displayed different patterns across study sites (i.e. significant interspecific trait difference x site interactions; Table S4). The frequent trait x site interactions arose because the numbers of traits that were significantly related to invasion growth rates decreased greatly towards high elevation (n = 9, 7 and 0 traits in the low, middle and high sites, respectively; Fig. 2), as did their effect sizes (F_2, 108_ = 26.288, P < 0.0001; Fig. S8 a&d). Although PCA axes captured species differences in multiple traits, their relationships with invasion growth rate also reduced greatly towards higher elevations. Specifically, PC1 only showed a significant relationship with invasion growth rates in the low and middle sites, PC2 only in the middle site, and PC3 only in the low site (Fig. 2). None of the three PCA axes was significantly related to invasion growth rates in the high site (Fig. 2). Similarly, among the 11 individual traits that were significantly related to relative fitness differences, seven of them had significantly different relationships between sites (Table S5). We found that ten traits were significantly related to relative fitness differences in the low site (Fig. 3 a&b), where species also displayed greater relative fitness differences (Fig. 1 d). The numbers of traits significantly related to relative fitness differences reduced to one and two in the middle and high sites, respectively (Fig. 3 a&b). The effect sizes of all traits also decreased greatly towards high elevations (F_2, 108_ = 35.53, P < 0.0001; Fig. S8 b&e). In analogue to individual traits, PC2 and PC3 were only significantly related to relative fitness differences at the low elevation (Fig. 3 a&b).

In contrast, the relationships between interspecific trait differences in both individual traits and PCA axes and niche overlap were relatively constant across the study sites. All seven traits except one (first flowering date, FFD) were similarly related to niche overlap between sites (Table S6). The results showed that species had greater niche overlap when they flowered at similar times at the middle site, but the relationship was reversed at the low site (Fig. 3 c&d). Consistently, we found similar numbers of traits that were significantly related to niche overlap across the elevation gradient (n = 5, 4 and 4 traits in the low, middle and high sites, respectively; Fig. 3 c&d) and the effect sizes of all traits were also similar between sites (F_2, 108_ = 0.969, P = 0.616; Fig. S8 c&f).

### How do the relationships between trait differences and coexistence differ between current vs novel competitors?

The relationships of both individual traits and PCA axes with invasion growth rates, relative fitness differences and niche overlap were very similar between current (lowland-lowland and highland-highland pairs) and novel (lowland-highland pairs) competitors. Among the 12 traits that were related to invasion growth rates, only one trait, first flowering date (FFD), had significantly different relationships for current versus novel competitors (Table S4). Similarly, all 11 traits that were related to relative fitness differences had similar relationships for novel and current competitors (Fig. 3 a&b; Table S5). Although our results showed that current competitors generally had smaller niche overlap than novel competitors (Fig. 1 e), only one (light interception, LI) of the seven traits that were related to niche overlap showed a significantly different effect for current vs novel pairs (i.e. significant interspecific trait difference x competitor origin interactions; Table S6); novel competitors with similar light interception abilities had greater niche overlap at the high site while the relationships were absent for current competitors (Fig. 3 c&d). The generally similar relationships between trait differences and coexistence for novel and current pairs were also supported by an analysis showing relatively consistent trait effect sizes between them (Fig. S8; invasion growth rate: F_2, 108_ = 0.722, P = 0.697; relative fitness differences: F_2, 108_ = 0.024, P = 0.988; niche overlap: F_2, 108_ = 1.649, P = 0.438), indicating that traits were very similarly related to both current and novel competition.

## Discussion

### Traits can predict outcomes of competition

Earlier work has shown the potential of using traits to inform our understanding of competition, and to forecast its impacts in plant communities (Kraft *et al*. 2015; Kunstler *et al*. 2016; Schleuning *et al*. 2020). But until now, whether traits are related to population-level outcomes of competition, and to what extent these relationships are generalisable under different climates and between current versus novel competitors, has rarely been tested. We found that traits associated with various ecological functions were correlated with the population-level outcomes of competition (i.e. invasion growth rates), implicating several processes as possibly mediating competition in this study system (Craine & Dybzinski 2013). In particular, the strong links between competition and traits associated with species’ ability to obtain light, such as light interception, plant height or leaf area, support a major role for light competition in mediating species’ interactions in these plant communities (Dybzinski & Tilman 2007; Vojtech *et al*. 2007; Violle *et al*. 2009; Williams *et al*. 2021). In addition, the significant effects of leaf nitrogen content, root traits and water use efficiency suggest that competition for soil water or nutrients was also an important driver of interactions in our experiment (Silvertown *et al*. 1999; Dybzinski & Tilman 2007). Nonetheless, it is important to note that these mechanisms of competition might be accentuated in experiments such as ours, where the relatively homogenous and stable environmental conditions limit the opportunities for niche differentiation that might exist in natural plant communities (e.g. environment heterogeneity and temporal fluctuations).

The fact that most traits related to invasion growth rates were also related to relative fitness differences, but rarely to niche overlap, suggests that the impacts of individual traits on competitive outcomes are mainly driven by how trait differences between species are linked to differences in their competitive ability (Kraft *et al*. 2015; Perez-Ramos *et al*. 2019). This result is not unexpected, since other studies have found that individual traits usually characterize species’ competitive ability, while species’ niche differences are more likely to be determined by multiple traits simultaneously (Kraft *et al*. 2015; Pérez-Ramos *et al*. 2019). In line with this, our multivariate analyses (Supplementary methods) revealed that certain trait combinations, in some cases, outperformed individual traits in influencing niche overlap (Fig. S11), though in some cases this was also true for invasion growth rates (Fig. S9) and relative fitness differences (Fig. S10), underlining the multidimensional nature of species coexistence (Clark *et al*. 2007; Kraft *et al*. 2015). Nevertheless, increasing the number of traits did not always result in stronger relationships between trait differences and coexistence. For example, the best (lowest-AICc) models for relative fitness differences included the combinations of five, one and four traits in the low, middle and high sites, respectively (Fig. S10). This suggests that particular combinations of traits are needed to explain species coexistence, and might also partly explain why PCA axes containing information on multiple traits were not always more strongly related to relative fitness differences or niche overlap than individual traits in our system.

### Traits are better predictors of competition under warmer, lower elevation sites

Functional traits were more strongly associated with the outcomes of competition at lower elevations, where the climate is warmer. This is in line with a recent study showing that plant height and specific leaf area are more strongly related to tree species‘ responses to neighbours in tropical forests and temperate rainforests than in taiga ecosystems (Kunstler *et al*. 2016). We suggest two reasons for the weaker effects of traits on competition under colder climate. First, the weakened effects of some traits on the outcomes of competition at higher elevations could be due to the shifts in limiting resources across the elevation gradient (e.g. Geng *et al*. 2017). For example, we found that traits characterising species’ abilities to obtain light were strongly related to their competitive ability at the lower but not high elevations, suggesting a greater role of competition for light under warmer climates (Walker *et al*. 2006; Hautier *et al*. 2018; Martin *et al*. 2020). Therefore, it is possible that traits not measured in this study could have significant effects on competition at higher elevations, such as those associated with frost resistance (Körner 2016). Further work would be needed to understand resource or non-resource-based mechanisms of competition and how this changes with elevation in order to identify traits that can capture variation in competition outcomes at high as well as at low elevation.

In addition to the possible shifts in limiting resources across the elevation gradient, our results also suggest that the effects of traits on the outcomes of competition appear to be weaker when species display more similar trait values. We found that, at higher elevations, species were less differentiated in functional trait space (Fig. S3) and many traits displayed smaller interspecific differences (Fig. S4), perhaps due to more limited growth overall. Furthermore, the weakened association between traits and the outcomes of competition was linked to a reduced ability of traits to capture relative fitness differences towards higher elevations. One possible reason for this could be that a cooler climate and shorter growing season constrain individual growth and trait expression, effectively equalizing species’ competitive abilities (Fig. S7 d).

In contrast to trait associations with relative fitness differences, trait associations with niche overlap were largely unchanged across the elevation gradient. One possible reason for these contrasting environmental dependencies may be the distinct trait architectures (i.e. the number of traits and their relative effects) underlying competitive ability versus niche overlap (Kraft *et al*. 2015; Pérez-Ramos *et al*. 2019), as discussed above, which might be worthy of further investigation. These results suggest that traits may better predict the outcomes of competition under warmer climate or in high-productivity ecosystems, and ongoing climatic warming might, therefore, tend to enhance their predictive ability.

### Traits predict competition equally well for current and novel competitors

Contrary to our hypothesis, we found that the effects of traits on coexistence were very similar for current and novel pairs of competitors. This result indicates that competition between current and novel competitors is mediated by similar processes, such as competition for light and soil water and nutrients, as discussed above. Nonetheless, we deliberately selected high and low-elevation species to have high overlap in functional trait space (see Methods; Fig. S3), such that current and novel pairs had similar magnitudes of trait differences (Fig. S4). Had we included, for example, only tall lowland and short highland species, which typically characterise the low and high-elevation communities, respectively, we might have found that height was a better predictor of novel competition, simply because novel competitors had greater differences in plant height than current competitors (analogous to the greater effects of traits on coexistence we have observed towards lower elevations and described in the previous section). Therefore, including lowland and highland species that had similar trait values allowed us to focus on the impacts of co-occurrence history per se on trait-competition associations, such as through competition-driven character displacement.

The fact that current (sympatric) competitors had smaller niche overlap but similar magnitudes of relative fitness differences compared to novel (allopatric) competitors (Fig. S7) suggests that the co-occurrence history between competing species may facilitate coexistence mainly via its impacts on niche overlap, potentially through the divergence of traits associated with species’ niche use (Germain *et al*. 2018b). Interestingly, we did not detect any significant changes to the effects of traits on niche overlap between current vs novel competitors. A possible reason is that co-occurrence history reduces niche overlap between current competitors through simultaneous but small shifts in the expression of multiple traits (Kraft *et al*. 2015; Germain *et al*. 2018b). In fact, species’ abilities to intercept light, the only trait that showed different impacts for current vs novel competitors, is likely to be determined by multiple traits, such as plant height and leaf economic traits. Although we measured traits associated with various plant functions, the possibility remains that co-occurrence history might reduce niche overlap through the displacement of traits not measured in this study. Apart from that, the generally consistent effects of traits on coexistence between current and novel competitors suggest that similar mechanisms govern their interactions.

### Implications for the impacts of changing interactions on plant communities

By linking functional traits to the processes determining the outcomes of competition under different climates, our study provides important insight into how altered species interactions might mediate the impacts of changing climate on plant communities. We found that the frequency of competitive exclusion increased towards lower elevations mainly due to amplified relative fitness differences, likely resulting from intensified competition for light (Sauter *et al*. 2021). This result suggests that climate warming may increase the chance of competitive exclusion of species that are weaker competitors for light, thereby reducing community diversity (Tylianakis *et al*. 2008). In addition, we observed greater niche overlap between novel competitors than between species that currently interact, and accordingly, a higher frequency of competitive exclusion in pairs of novel competitors (Fig. S7). This provides further evidence that novel competitors can potentially exert greater competitive impacts than members of the local community (Alexander *et al*. 2015). Our experiment adds to a growing body of studies showing that climate change can systematically alter the impacts of competition on community members and reinforces the importance of considering the impacts of changing interactions for understanding and forecasting the responses of plant communities to climate change (Tylianakis *et al*. 2008; Zhang *et al*. 2015; Vandvik *et al*. 2020).

Although we found an increased frequency of stable coexistence towards higher elevations, the intensity of interspecific competition per se, measured as competition-induced suppression of population growth, did not systematically decrease with increasing elevation in our experiment (Fig. S5), contrasting conventional hypotheses (e.g. the stress-gradient hypothesis) (Callaway 1998). Instead, our results showed that it was the smaller differences in competitive ability between interacting species (i.e. equalized relative fitness differences) that contributed to the greater chance of coexistence at higher elevations. Importantly, this does not necessarily translate into a reduced importance of competition towards high elevations or cold climate (Freckleton *et al*. 2009; Louthan *et al*. 2015; Lyu & Alexander 2022). Therefore, more empirical studies employing such theoretically motivated approaches are needed to gain insights into the varying roles of species interactions in structuring ecological communities across environmental gradients (Chesson & Huntly 1997; Louthan *et al*. 2015; Germain *et al*. 2018a).

### Caveats and future directions

Although our study demonstrates the usefulness of species’ traits for predicting the outcomes of current and novel competition under different climate, the traits we measured only explained small proportions of variance in invasion growth rates (e.g. light interception had the greatest predictive power but only accounted for 11% of the variance; Table S4). In addition to the possible omission of other important traits, as discussed above, we suggest two other possible reasons for this. First, individual-level trait plasticity in response to competition (e.g. a focal plant can have different reproductive heights depending on its competitors) might affect population growth under competition (Bennett *et al*. 2016).

Nevertheless, we found that species’ differences in reproductive height measured on each species pair were not more closely related to invasion growth rates than those based on the grand species-level mean (Fig. S12). Secondly, we showed that coexistence may be mediated by multiple traits simultaneously, which might limit the variance in competition outcomes explained by individual traits (Clark *et al*. 2007; Kraft *et al*. 2015; Pérez-Ramos *et al*. 2019; Pistón *et al*. 2019). Future enquiries into these aspects in various systems are needed to improve our understanding of, and ability to predict, the impacts of competition on plant communities under changing climate.

Functional traits are already being used to predict community composition in response to climate change (Isabelle *et al*. 2014; Cadotte *et al*. 2015; Schleuning *et al*. 2020) and incorporated into mechanistic models to inform projections of species range shifts following climate changes (MacLean & Beissinger 2017; Briscoe *et al*. 2019; Guisan *et al*. 2019). Our study provides a test of several key assumptions underlying these trait-based approaches by establishing the links between functional traits and the outcome of competition and the processes determining these outcomes under different climatic conditions and for current and novel interactions (Alexander *et al*. 2016). For the first time to our knowledge, our study demonstrates that traits can predict the outcome of interactions following range shifts among species that do not yet interact. Additionally, we show that the predictive ability of traits depends greatly on environmental conditions, suggesting that prior information may be needed on the particular limiting factors and relevant traits that can be expected under altered climate in order to reliably predict species’ responses to changing species interactions. Overall, our study contributes to developing a theoretically justified trait-based approach to forecasting the responses of plant populations and communities to environmental changes (McGill *et al*. 2006; Shipley *et al*. 2016).

## Acknowledgements

We thank members of the Plant Ecology Group at ETH Zürich for assistance with field and lab work, in particular Loïc Liberati, Tim Murray, Yan Hess, Jessica Joaquim, Megan Stamp, Annette Altermatt, and Camille Brioschi. We thank the Commune de Bex and Jean-Louis Putallaz for the access to the field sites. We thank the Stable Isotope Laboratory of Grassland Sciences Group at ETH Zürich for help with leaf chemical and carbon isotope analyses. We also thank anonymous reviewers for their comments and suggestions that helped improved the manuscript. S.L. also thanks the Chinese Scholarship Council for financial support (No.201706100184). J.M.A. received funding from the European Union’s Horizon 2020 research and innovation programme under grant agreement No. 678841.

## Authorship

J.M.A. designed the field experiment. S.L. and J.M.A. collected the data. S.L. conducted the data analyses and wrote the manuscript with input from J.M.A.

## References

Adams, T.P., Purves, D.W. & Pacala, S.W. (2007). Understanding height-structured competition in forests: is there an R* for light? Proc. R. Soc. B-Biol. Sci., 274, 3039–3047.

Adler, P.B., Fajardo, A., Kleinhesselink, A.R. & Kraft, N.J.B. (2013). Trait-based tests of coexistence mechanisms. Ecology Letters, 16, 1294–1306.

Alexander, J.M., Diez, J.M., Hart, S.P. & Levine, J.M. (2016). When Climate Reshuffles Competitors: A Call for Experimental Macroecology. Trends in Ecology & Evolution, 31, 831–841.

Alexander, J.M., Diez, J.M. & Levine, J.M. (2015). Novel competitors shape species’ responses to climate change. Nature, 525, 515–518.

Arnold, S.J. (1983). Morphology, Performance and Fitness. American Zoologist, 23, 347–361.

Bennett, J.A., Riibak, K., Tamme, R., Lewis, R.J. & Pärtel, M. (2016). The reciprocal relationship between competition and intraspecific trait variation. Journal of Ecology, 104, 1410–1420.

Bjorkman, A.D., Myers-Smith, I.H., Elmendorf, S.C., Normand, S., Rüger, N., Beck, P.S.A. et al. (2018). Plant functional trait change across a warming tundra biome. Nature, 562, 57–62.

Blackford, C., Germain, R.M. & Gilbert, B. (2020). Species Differences in Phenology Shape Coexistence. *The American Naturalist*, E000–E000.

Borges, I.L., Forsyth, L.Z., Start, D. & Gilbert, B. (2019). Abiotic heterogeneity underlies trait-based competition and assembly. Journal of Ecology, 107, 747–756.

Briscoe, N.J., Elith, J., Salguero-Gómez, R., Lahoz-Monfort, J.J., Camac, J.S., Giljohann, K.M. et al. (2019). Forecasting species range dynamics with process-explicit models: matching methods to applications. Ecology Letters, 0.

Cadotte, M.W., Arnillas, C.A., Livingstone, S.W. & Yasui, S.-L.E. (2015). Predicting communities from functional traits. Trends in Ecology & Evolution, 30, 510–511.

Callaway, R.M. (1998). Competition and Facilitation on Elevation Gradients in Subalpine Forests of the Northern Rocky Mountains, USA. Oikos, 82, 561–573.

Carroll, I.T., Cardinale, B.J. & Nisbet, R.M. (2011). Niche and fitness differences relate the maintenance of diversity to ecosystem function. 92, 1157–1165.

Chesson, P. (2000). Mechanisms of Maintenance of Species Diversity. Annual Review of Ecology and Systematics, 31, 343–366.

Chesson, P. & Huntly, N. (1997). The Roles of Harsh and Fluctuating Conditions in the Dynamics of Ecological Communities. The American Naturalist, 150, 519–553.

Clark, J.S., Dietze, M., Chakraborty, S., Agarwal, P.K., Ibanez, I., LaDeau, S. et al. (2007). Resolving the biodiversity paradox. Ecology Letters, 10, 647–659.

Collins, C.G., Elmendorf, S.C., Smith, J.G., Shoemaker, L., Szojka, M., Swift, M. et al. (2022). Global change re-structures alpine plant communities through interacting abiotic and biotic effects. Ecology Letters, 25, 1813–1826.

Copeland, S.M. & Harrison, S.P. (2017). Community traits affect plant-plant interactions across climatic gradients. Oikos, 126, 296–305.

Craine, J.M. & Dybzinski, R. (2013). Mechanisms of plant competition for nutrients, water and light. Functional Ecology, 27, 833–840.

Descombes, P., Pitteloud, C., Glauser, G., Defossez, E., Kergunteuil, A., Allard, P.M. et al. (2020). Novel trophic interactions under climate change promote alpine plant coexistence. Science, 370, 1469–1473.

Díaz, S., Kattge, J., Cornelissen, J.H., Wright, I.J., Lavorel, S., Dray, S. et al. (2016). The global spectrum of plant form and function. Nature, 529, 167–171.

Dybzinski, R. & Tilman, D. (2007). Resource use patterns predict long-term outcomes of plant competition for nutrients and light. The American Naturalist, 170, 305–318.

Ellner, S.P., Childs, D.Z. & Rees, M. (2016). Data-Driven Modelling of Structured Populations : A Practical Guide to the Integral Projection Model. Cham : Springer.

Elmendorf, S.C., Henry, G.H.R., Hollister, R.D., Fosaa, A.M., Gould, W.A., Hermanutz, L. et al. (2015). Experiment, monitoring, and gradient methods used to infer climate change effects on plant communities yield consistent patterns. Proceedings of the National Academy of Sciences, 112, 448–452.

Freckleton, R.P., Watkinson, A.R. & Rees, M. (2009). Measuring the importance of competition in plant communities. Journal of Ecology, 97, 379–384.

Geng, Y., Baumann, F., Song, C., Zhang, M., Shi, Y., Kühn, P. et al. (2017). Increasing temperature reduces the coupling between available nitrogen and phosphorus in soils of Chinese grasslands. Scientific Reports, 7, 43524.

Germain, R.M., Mayfield, M.M. & Gilbert, B. (2018a). The “filtering” metaphor revisited: competition and environment jointly structure invasibility and coexistence. Biology Letters, 14, 20180460.

Germain, R.M., Williams, J.L., Schluter, D. & Angert, A.L. (2018b). Moving Character Displacement beyond Characters Using Contemporary Coexistence Theory. Trends in Ecology & Evolution, 33, 74–84.

Gilman, S.E., Urban, M.C., Tewksbury, J., Gilchrist, G.W. & Holt, R.D. (2010). A framework for community interactions under climate change. Trends in Ecology & Evolution, 25, 325–331.

Godoy, O. & Levine, J.M. (2014). Phenology effects on invasion success: insights from coupling field experiments to coexistence theory. Ecology, 95, 726–736.

Grainger, T.N., Levine, J.M. & Gilbert, B. (2019). The Invasion Criterion: A Common Currency for Ecological Research. Trends in Ecology & Evolution, 34, 925–935.

Guisan, A., Mod, H.K., Scherrer, D., Münkemüller, T., Pottier, J., Alexander, J.M. et al. (2019). Scaling the linkage between environmental niches and functional traits for improved spatial predictions of biological communities. Global Ecology and Biogeography, 28, 1384–1392.

Hautier, Y., Vojtech, E. & Hector, A. (2018). The importance of competition for light depends on productivity and disturbance. Ecology and Evolution, 8, 10655–10661.

Isabelle, B., Damien, G. & Wilfried, T. (2014). FATE-HD: a spatially and temporally explicit integrated model for predicting vegetation structure and diversity at regional scale. Glob Chang Biol, 20, 2368–2378.

Kleyer, M., Bekker, R.M., Knevel, I.C., Bakker, J.P., Thompson, K., Sonnenschein, M. et al. (2008). The LEDA Traitbase: a database of life-history traits of the Northwest European flora. Journal of Ecology, 96, 1266–1274.

Körner, C. (2016). Plant adaptation to cold climates. F1000Res, 5, F1000 Faculty Rev-2769.

Kraft, N.J.B., Godoy, O. & Levine, J.M. (2015). Plant functional traits and the multidimensional nature of species coexistence. P Natl Acad Sci USA, 112, 797–802.

Kunstler, G., Falster, D., Coomes, D.A., Hui, F., Kooyman, R.M., Laughlin, D.C. et al. (2016). Plant functional traits have globally consistent effects on competition. Nature, 529, 204–207.

Kunstler, G., Lavergne, S., Courbaud, B., Thuiller, W., Vieilledent, G., Zimmermann, N.E. et al. (2012). Competitive interactions between forest trees are driven by species’ trait hierarchy, not phylogenetic or functional similarity: implications for forest community assembly. Ecology Letters, 15, 831–840.

Li, Y., Jiang, Y., Zhao, K., Chen, Y., Wei, W., Shipley, B. et al. (2022). Exploring trait– performance relationships of tree seedlings along experimentally manipulated light and water gradients. Ecology, 103, e3703.

Louthan, A.M., Doak, D.F. & Angert, A.L. (2015). Where and When do Species Interactions Set Range Limits? Trends in Ecology & Evolution, 30, 780–792.

Lyu, S. & Alexander, J.M. (2022). Competition contributes to both warm and cool range edges. Nature Communications, 13, 2502.

Lyu, S. & Alexander, J.M. (2023). Compensatory responses of vital rates attenuate impacts of competition on population growth and promote coexistence. Ecology Letters, 26, 437–447.

Lyu, S., Liu, X., Venail, P. & Zhou, S. (2017). Functional dissimilarity, not phylogenetic relatedness, determines interspecific interactions among plants in the Tibetan alpine meadows. Oikos, 126, 381–388.

MacLean, S.A. & Beissinger, S.R. (2017). Species’ traits as predictors of range shifts under contemporary climate change: A review and meta-analysis. Global Change Biology, 23, 4094–4105.

Martin, J.T., Pederson, G.T., Woodhouse, C.A., Cook, E.R., McCabe, G.J., Anchukaitis, K.J. et al. (2020). Increased drought severity tracks warming in the United States’ largest river basin. Proceedings of the National Academy of Sciences, 117, 11328–11336.

Mathakutha, R., Steyn, C., le Roux, P.C., Blom, I.J., Chown, S.L., Daru, B.H. et al. (2019). Invasive species differ in key functional traits from native and non-invasive alien plant species. Journal of Vegetation Science, 30, 994–1006.

Matías, L., Godoy, O., Gómez-Aparicio, L. & Pérez-Ramos, I.M. (2018). An experimental extreme drought reduces the likelihood of species to coexist despite increasing intransitivity in competitive networks. Journal of Ecology, 106, 826–837.

McGill, B.J., Enquist, B.J., Weiher, E. & Westoby, M. (2006). Rebuilding community ecology from functional traits. Trends Ecol. Evol., 21, 178–185.

Narwani, A., Alexandrou, M.A., Oakley, T.H., Carroll, I.T. & Cardinale, B.J. (2013). Experimental evidence that evolutionary relatedness does not affect the ecological mechanisms of coexistence in freshwater green algae. Ecology Letters, 16, 1373–1381.

Nomoto, H.A. & Alexander, J.M. (2021). Drivers of local extinction risk in alpine plants under warming climate. Ecol. Lett., 24, 1157–1166.

Perez-Harguindeguy, N., Diaz, S., Garnier, E., Lavorel, S., Poorter, H., Jaureguiberry, P. et al. (2013). New handbook for standardised measurement of plant functional traits worldwide. Australian Journal of Botany, 61, 167–234.

Perez-Ramos, I.M., Matias, L., Gomez Aparicio, L. & Godoy, O. (2019). Functional traits and phenotypic plasticity modulate species coexistence across contrasting environments. bioRxiv, 539619.

Pérez-Ramos, I.M., Matías, L., Gómez-Aparicio, L. & Godoy, Ó. (2019). Functional traits and phenotypic plasticity modulate species coexistence across contrasting climatic conditions. Nature Commun., 10, 2555.

Pistón, N., de Bello, F., Dias, A.T.C., Götzenberger, L., Rosado, B.H.P., de Mattos, E.A. et al. (2019). Multidimensional ecological analyses demonstrate how interactions between functional traits shape fitness and life history strategies. Journal of Ecology, 107, 2317–2328.

R Core Team (2020). R: A language and environment for statistical computing. R Foundation for Statistical Computing Vienna, Austria.

Randin, C.F., Engler, R., Normand, S., Zappa, M., Zimmermann, N.E., Pearman, P.B. et al. (2009). Climate change and plant distribution: local models predict high-elevation persistence. Glob. Chang. Biol., 15, 1557–1569.

Rumpf, S.B., Hülber, K., Klonner, G., Moser, D., Schütz, M., Wessely, J. et al. (2018). Range dynamics of mountain plants decrease with elevation. Proceedings of the National Academy of Sciences, 115, 1848–1853.

Sakarchi, J. & Germain, R.M. (2023). The Evolution of Competitive Ability. The American Naturalist, 201, 1–15.

Sauter, F., Albrecht, H., Kollmann, J. & Lang, M. (2021). Competition components along productivity gradients -revisiting a classic dispute in ecology. Oikos, 130, 1326–1334.

Scherrer, D., Vitasse, Y., Guisan, A., Wohlgemuth, T. & Lischke, H. (2020). Competition and demography rather than dispersal limitation slow down upward shifts of trees’ upper elevation limits in the Alps. Journal of Ecology, 108, 2416–2430.

Schleuning, M., Neuschulz, E.L., Albrecht, J., Bender, I.M.A., Bowler, D.E., Dehling, D.M. et al. (2020). Trait-Based Assessments of Climate-Change Impacts on Interacting Species. Trends Ecol. Evol., 35, 319–328.

Sherry, R.A., Zhou, X., Gu, S., Arnone, J.A., Schimel, D.S., Verburg, P.S. et al. (2007). Divergence of reproductive phenology under climate warming. Proceedings of the National Academy of Sciences, 104, 198–202.

Shipley, B., De Bello, F., Cornelissen, J.H.C., Laliberté, E., Laughlin, D.C. & Reich, P.B. (2016). Reinforcing loose foundation stones in trait-based plant ecology. Oecologia, 180, 923–931.

Silvertown, J., Dodd, M.E., Gowing, D.J.G. & Mountford, J.O. (1999). Hydrologically defined niches reveal a basis for species richness in plant communities. Nature, 400, 61–63.

Spaak, J.W. & De Laender, F. (2020). Intuitive and broadly applicable definitions of niche and fitness differences. Ecology Letters, 23, 1117–1128.

Suttle, K.B., Thomsen, M.A. & Power, M.E. (2007). Species interactions reverse grassland responses to changing climate. Science, 315, 640–642.

Tilman, D. (1982). Resource competition and community structure. Princeton University Press, Princeton, N.J.

Tylianakis, J.M., Didham, R.K., Bascompte, J. & Wardle, D.A. (2008). Global change and species interactions in terrestrial ecosystems. Ecol Lett, 11, 1351–1363.

Van Dyke, M.N., Levine, J.M. & Kraft, N.J.B. (2022). Small rainfall changes drive substantial changes in plant coexistence. Nature.

van Kleunen, M., Weber, E. & Fischer, M. (2009). A Meta-Analysis of Trait Differences Between Invasive and Non-Invasive Plant Species. Ecology letters, 13, 235–245.

Vandvik, V., Skarpaas, O., Klanderud, K., Telford, R.J., Halbritter, A.H. & Goldberg, D.E. (2020). Biotic rescaling reveals importance of species interactions for variation in biodiversity responses to climate change. Proceedings of the National Academy of Sciences, 117, 22858–22865.

Violle, C., Garnier, E., Lecoeur, J., Roumet, C., Podeur, C., Blanchard, A. et al. (2009). Competition, traits and resource depletion in plant communities. Oecologia, 160, 747–755.

Vitasse, Y., Ursenbacher, S., Klein, G., Bohnenstengel, T., Chittaro, Y., Delestrade, A. et al. (2021). Phenological and elevational shifts of plants, animals and fungi under climate change in the European Alps. Biological Reviews, 96, 1816–1835.

Vojtech, E., Turnbull, L.A. & Hector, A. (2007). Differences in Light Interception in Grass Monocultures Predict Short-Term Competitive Outcomes under Productive Conditions. PLOS ONE, 2, e499.

Walker, M.D., Wahren, C.H., Hollister, R.D., Henry, G.H.R., Ahlquist, L.E., Alatalo, J.M. et al. (2006). Plant community responses to experimental warming across the tundra biome. Proceedings of the National Academy of Sciences, 103, 1342–1346.

Walther, G.-R., Post, E., Convey, P., Menzel, A., Parmesan, C., Beebee, T.J.C. et al. (2002). Ecological responses to recent climate change. Nature, 416, 389–395.

Wei, B., Zhang, D., Wang, G., Liu, Y., Li, Q., Zheng, Z. et al. Experimental warming altered plant functional traits and their coordination in a permafrost ecosystem. New Phytologist, n/a.

Williams, L.J., Butler, E.E., Cavender-Bares, J., Stefanski, A., Rice, K.E., Messier, C. et al. (2021). Enhanced light interception and light use efficiency explain overyielding in young tree communities. Ecology Letters, 24, 996–1006.

Wright, I.J., Reich, P.B., Westoby, M., Ackerly, D.D., Baruch, Z., Bongers, F. et al. (2004). The worldwide leaf economics spectrum. Nature, 428, 821–827.

Yang, X., Gómez-Aparicio, L., Lortie, C.J., Verdú, M., Cavieres, L.A., Huang, Z. et al. (2022). Net plant interactions are highly variable and weakly dependent on climate at the global scale. Ecology Letters, 25, 1580–1593.

Zhang, J., Huang, S. & He, F. (2015). Half-century evidence from western Canada shows forest dynamics are primarily driven by competition followed by climate. Proceedings of the National Academy of Sciences, 112, 4009–4014.

Zhang, Z. & van Kleunen, M. (2019). Common alien plants are more competitive than rare natives but not than common natives. Ecol. Lett., 22, 1378–1386.

Zuppinger-Dingley, D., Schmid, B., Petermann, J.S., Yadav, V., De Deyn, G.B. & Flynn, D.F.B. (2014). Selection for niche differentiation in plant communities increases biodiversity effects. Nature, 515, 108–111.

