## Supplementary information for "Functional traits predict outcomes of current and novel competition under warmer climate"

### Supplementary Information for the manuscript: Functional traits predict outcomes of current and novel interactions under warmer climate

Shengman Lyu1[1,2*], Jake M. Alexander[1]

[1] Institute of Integrative Biology, ETH Zürich, 8092 Zürich, Switzerland

[2] Department of Ecology and Evolution, University of Lausanne, 1015 Lausanne, Switzerland

#### Supplementary methods

*Multivariate analyses*

We performed multivariate analyses to test whether multiple traits in certain combinations that were not captured by PCA axes affect coexistence. We first calculated the interspecific difference of multiple traits for combinations of traits including two to 15 traits. Interspecific differences of multiple traits were calculated as the Euclidean distance in the multidimensional space defined by a set of traits (Petchey & Gaston 2006). Given the enormous number of all possible combinations (n = 32768 in total), we randomly drew 5% (n = 1638). For each trait combination, we tested the effects of interspecific differences of multiple traits on invasion growth rates, relative fitness differences and niche overlap in each site separately and excluded the origin of competitors as a covariable from the models as it never affected the effects of individual traits (Table S4). We used the Akaike information criterion corrected for small sample size (AICc) to select the best-fit models.

*Supplementary references*

Petchey, O.L. & Gaston, K.J. (2006). Functional diversity: back to basics and looking forward. *Ecology Letters*, 9, 741-758.

**Table S1**. Species included in this study. The elevation range is defined as the 10th and 90th percentile of a species' elevation distribution in the study area.

| **Species** | **Family** | **Functional group** | **Origin of elevation** | **Growth form** | **Elevation range (m, mean, lower-upper)** | **Seed supplier** |
|---|---|---|---|---|---|---|
| *Bromus erectus* | Poaceae | Grass | Lowland | Perennial | 971, 598-1351 | Otto Hauenstein Samen |
| *Crepis biennis* | Asteraceae | Forb | Lowland | Biennial | 1007, 764-1299 | Otto Hauenstein Samen |
| *Daucus carota* | Apiaceae | Forb | Lowland | Biennial | 1063, 683-1429 | UFA SAMEN |
| *Medicago lupulina* | Fabaceae | Legume | Lowland | Perennial | 1035, 653-1408 | Otto Hauenstein Samen |
| *Plantago lanceolata* | Plantaginaceae | Forb | Lowland | Perennial | 1169, 629-1657 | UFA SAMEN |
| *Poa trivialis* | Poaceae | Grass | Lowland | Perennial | 980, 527-1390 | UFA SAMEN |
| *Salvia pratensis* | Lamiaceae | Forb | Lowland | Perennial | 782, 539-1069 | Otto Hauenstein Samen |
| *Anthylis vulneraria ssp. alpestris* | Fabaceae | Legume | Highland | Perennial | 1848, 1341-2217 | Kaertner Saatbau |
| *Arnica montana* | Asteraceae | Forb | Highland | Perennial | 1906, 1622-2091 | Jellito |
| *Aster alpinus* | Asteraceae | Forb | Highland | Perennial | 2108, 2002-2236 | Jellito |
| *Plantago alpina* | Plantaginaceae | Forb | Highland | Perennial | 1916, 1581-2193 | Schutz Filisur |
| *Poa alpina* | Poaceae | Grass | Highland | Perennial | 2045, 1674-2458 | Kaertner Saatbau |
| *Sesleria caerulea* | Poaceae | Grass | Highland | Perennial | 2013, 1652-2371 | Jellito |
| *Trifolium badium* | Fabaceae | Legume | Highland | Perennial | 1943, 1640-2253 | Schutz Filisur |

**Table S2**. Correlations (Pearson's *r*) of functional traits between species. Correlations between traits are calculated by pooling trait data across the three study sites together, and only species mean are used. Strong correlations between traits with the absolute values of Pearson's *r* greater than 0.5 are highlighted in boldface. See Table 1 for trait abbreviations.

| | RH | VH | RMF | LA | LDMC | SLA | LN | LC | SM | FRP | FRD | SRL | FFD | δ13C |
|---|---|---|---|---|---|---|---|---|---|---|---|---|---|---|
| VH | 0.487 | | | | | | | | | | | | | |
| RMF | -0.321 | -0.123 | | | | | | | | | | | | |
| LA | 0.435 | 0.206 | 0.106 | | | | | | | | | | | |
| LDMC | 0.082 | 0.224 | -0.189 | **-0.536** | | | | | | | | | | |
| SLA | -0.133 | **-0.631** | -0.025 | -0.134 | -0.191 | | | | | | | | | |
| LN | -0.329 | **-0.612** | 0.046 | -0.098 | -0.254 | **0.724** | | | | | | | | |
| LC | -0.428 | -0.064 | 0.303 | -0.176 | 0.394 | 0.121 | 0.264 | | | | | | | |
| SM | 0.297 | 0.461 | 0.027 | 0.382 | -0.080 | -0.185 | -0.055 | -0.019 | | | | | | |
| FRP | -0.023 | 0.173 | -0.249 | -0.238 | 0.340 | 0.284 | 0.035 | 0.043 | 0.104 | | | | | |
| FRD | -0.028 | **0.544** | 0.118 | 0.005 | **0.549** | **-0.526** | -0.387 | **0.529** | 0.237 | 0.069 | | | | |
| SRL | 0.250 | -0.192 | -0.372 | -0.400 | 0.401 | 0.116 | 0.016 | -0.064 | -0.496 | -0.165 | -0.213 | | | |
| FFD | 0.020 | -0.323 | -0.306 | -0.002 | -0.039 | 0.118 | -0.123 | -0.250 | -0.013 | -0.075 | -0.183 | 0.100 | | |
| δ13C | -0.122 | -0.229 | 0.296 | 0.411 | **-0.675** | 0.188 | 0.335 | -0.152 | 0.435 | -0.306 | -0.381 | **-0.540** | 0.176 | |
| LI | -0.433 | -0.235 | 0.051 | **-0.564** | 0.225 | 0.209 | 0.064 | 0.318 | -0.324 | 0.277 | -0.018 | 0.214 | 0.001 | -0.293 |

**Table S3.** interspecific trait differences (|$Trait_{focal}$ – $Trait_{competitor}$|) vary between sites (low, middle and high) and competitor origin (current and novel competitors). The effects were tested using mixed-effects models for each trait separately, with interspecific trait differences of species pairs (n=250) as response variables and site (low, middle or high) and origin of competitors (current or novel) and their interaction as fixed effects and the identity of focal (13) and competitor (12) species as random effects. P values smaller than 0.05 are in boldface. See Table 1 for trait abbreviations.

| **Trait** | **Site** | | **Origin** | | **Site:Origin** | |
|---|---|---|---|---|---|---|
| | d.f. = 2 | | 1 | | 2 | |
| | *F* | *P* | *F* | *P* | *F* | *P* |
| RH | 1.881 | 0.390 | 3.631 | 0.057 | 0.093 | 0.954 |
| VH | 3.469 | 0.177 | 4.095 | **0.043** | 1.258 | 0.533 |
| RMF | 26.954 | **<0.001** | 0.042 | 0.838 | 2.316 | 0.314 |
| LA | 2.504 | 0.286 | 7.911 | **0.005** | 0.101 | 0.951 |
| LDMC | 1.615 | 0.446 | 3.151 | 0.076 | 0.336 | 0.845 |
| SLA | 2.811 | 0.245 | 0.003 | 0.956 | 0.544 | 0.762 |
| LN | 0.865 | 0.649 | 3.326 | 0.068 | 0.799 | 0.671 |
| LC | 8.839 | **0.012** | 7.395 | **0.007** | 0.476 | 0.788 |
| FRP | 57.069 | **<0.001** | 0.659 | 0.417 | 0.047 | 0.977 |
| FRD | 1.588 | 0.452 | 6.675 | **0.010** | 3.435 | 0.180 |
| SRL | 14.870 | **0.001** | 0.407 | 0.523 | 2.280 | 0.320 |
| SM | 3.116 | 0.211 | 8.630 | **0.003** | 0.150 | 0.928 |
| $\delta^{13}C$ | 15.462 | **<0.001** | 3.616 | 0.057 | 1.238 | 0.538 |
| FFD | 66.672 | **<0.001** | 0.296 | 0.586 | 0.573 | 0.751 |
| Light | 1.011 | 0.603 | 1.059 | 0.304 | 0.490 | 0.783 |
| PC1 | 4.685 | 0.096 | 2.716 | 0.099 | 0.883 | 0.643 |
| PC2 | 7.150 | **0.028** | 6.399 | **0.011** | 0.042 | 0.979 |
| PC3 | 6.590 | **0.037** | 1.970 | 0.160 | 3.027 | 0.220 |

**Table S4**. Effects of interspecific trait difference (TD; $Trait_{focal} – Trait_{competitor}$), site (low, middle or high), the origin of competitors (current or novel) and the interactions between variables on invasion growth rate (*n* = 250 across all species and sites). The effects were tested using mixed-effects models for each trait, with focal (*n* = 13) and competitor (*n* = 12) species as random effects. P values smaller than 0.05 are in boldface. See Table 1 for trait abbreviations.

| **Trait** | **TD** | | **Site** | | **Origin** | | **TD:Site** | | **TD:Origin** | | **$R^2_{marg}$** | **$R^2_{cond}$** |
|---|---|---|---|---|---|---|---|---|---|---|---|---|
| | *d.f.* = 1 | | 2 | | 1 | | 2 | | 1 | | | |
| | *F* | *P* | *F* | *P* | *F* | *P* | *F* | *P* | *F* | *P* | | |
| RH | 7.345 | **0.007** | 4.542 | 0.103 | 2.764 | 0.096 | 40.411 | **<0.001** | 1.050 | 0.305 | 0.099 | 0.693 |
| VH | 8.201 | **0.004** | 4.898 | 0.086 | 2.416 | 0.120 | 31.768 | **<0.001** | 0.173 | 0.678 | 0.098 | 0.707 |
| RMF | 2.001 | 0.157 | 4.066 | 0.131 | 2.426 | 0.119 | 6.248 | **0.044** | 0.018 | 0.893 | 0.022 | 0.647 |
| LA | 5.978 | **0.014** | 3.713 | 0.156 | 2.690 | 0.101 | 21.299 | **<0.001** | 0.117 | 0.732 | 0.067 | 0.657 |
| LDMC | 1.076 | 0.300 | 3.186 | 0.203 | 2.781 | 0.095 | 7.330 | **0.026** | 0.000 | 0.994 | 0.028 | 0.631 |
| SLA | 0.243 | 0.622 | 4.022 | 0.134 | 2.294 | 0.130 | 17.362 | **<0.001** | 0.001 | 0.977 | 0.038 | 0.669 |
| LN | 0.003 | 0.954 | 3.902 | 0.142 | 2.374 | 0.123 | 19.995 | **<0.001** | 0.235 | 0.628 | 0.038 | 0.671 |
| LC | 0.000 | 0.984 | 3.713 | 0.156 | 2.490 | 0.115 | 10.390 | **0.006** | 0.979 | 0.323 | 0.028 | 0.648 |
| FRP | 4.531 | **0.033** | 4.031 | 0.133 | 2.185 | 0.139 | 1.690 | 0.430 | 2.189 | 0.139 | 0.038 | 0.638 |
| FRD | 0.695 | 0.404 | 3.632 | 0.163 | 2.316 | 0.128 | 9.635 | **0.008** | 0.745 | 0.388 | 0.027 | 0.657 |
| SRL | 0.184 | 0.668 | 3.765 | 0.152 | 2.689 | 0.101 | 5.659 | 0.059 | 0.625 | 0.429 | 0.018 | 0.646 |
| SM | 1.561 | 0.212 | 3.630 | 0.163 | 2.328 | 0.127 | 11.498 | **0.003** | 0.021 | 0.885 | 0.045 | 0.640 |
| $\delta^{13}C$ | 0.049 | 0.825 | 3.252 | 0.197 | 2.541 | 0.111 | 3.053 | 0.217 | 0.374 | 0.541 | 0.017 | 0.629 |
| FFD | 0.022 | 0.883 | 3.283 | 0.194 | 2.427 | 0.119 | 13.815 | **0.001** | 4.550 | **0.033** | 0.042 | 0.662 |
| LI | 19.420 | **<0.001** | 5.002 | 0.082 | 2.326 | 0.127 | 30.108 | **<0.001** | 0.039 | 0.843 | 0.110 | 0.708 |
| PC1 | 7.013 | **0.008** | 5.156 | 0.076 | 2.456 | 0.117 | 38.423 | **<0.001** | 0.035 | 0.851 | 0.085 | 0.705 |
| PC2 | 4.898 | **0.027** | 3.061 | 0.216 | 2.740 | 0.098 | 7.665 | **0.022** | 0.104 | 0.747 | 0.046 | 0.627 |
| PC3 | 3.574 | 0.059 | 4.314 | 0.116 | 2.491 | 0.115 | 8.414 | **0.015** | 1.222 | 0.269 | 0.037 | 0.655 |

**Table S5.** Effects of interspecific trait differences (TD; $Trait_{focal}$ – $Trait_{competitor}$), site (low, middle or high), the origin of competitors (current or novel) and their interactions on relative fitness differences (*n* = 93 across all species and sites). The effects were tested using mixed-effects models for each trait, with species as random effects. P values smaller than 0.05 are in boldface. See Table 1 for trait abbreviations.

| Trait | TD | | Site | | Origin | | TD:Site | | TD:Origin | | $R^2_{marg}$ | $R^2_{cond}$ |
|---|---|---|---|---|---|---|---|---|---|---|---|---|
| | *d.f.* = 1 | | 2 | | 1 | | 2 | | 1 | | | |
| | *F* | *P* | *F* | *P* | *F* | *P* | *F* | *P* | *F* | *P* | | |
| RH | 1.379 | 0.240 | 0.350 | 0.839 | 0.189 | 0.664 | 24.965 | **<0.001** | 0.012 | 0.912 | 0.143 | 0.593 |
| VH | 0.698 | 0.404 | 0.216 | 0.898 | 0.404 | 0.525 | 12.390 | **0.002** | 0.114 | 0.736 | 0.081 | 0.578 |
| RMF | 7.799 | **0.005** | 0.475 | 0.789 | 0.241 | 0.624 | 8.147 | **0.017** | 0.844 | 0.358 | 0.115 | 0.543 |
| LA | 1.945 | 0.163 | 0.313 | 0.855 | 0.328 | 0.567 | 8.170 | **0.017** | 0.000 | 0.996 | 0.072 | 0.522 |
| LDMC | 5.628 | **0.018** | 0.270 | 0.874 | 0.244 | 0.621 | 1.876 | 0.391 | 0.010 | 0.922 | 0.085 | 0.482 |
| SLA | 2.108 | 0.147 | 0.297 | 0.862 | 0.174 | 0.677 | 0.936 | 0.626 | 0.236 | 0.627 | 0.048 | 0.524 |
| LN | 2.893 | 0.089 | 0.360 | 0.835 | 0.338 | 0.561 | 1.459 | 0.482 | 0.129 | 0.720 | 0.051 | 0.492 |
| LC | 4.541 | **0.033** | 0.321 | 0.852 | 0.395 | 0.530 | 15.833 | **<0.001** | 0.054 | 0.815 | 0.129 | 0.552 |
| FRP | 0.047 | 0.828 | 0.354 | 0.838 | 0.376 | 0.540 | 1.413 | 0.493 | 0.005 | 0.944 | 0.013 | 0.490 |
| FRD | 4.578 | **0.032** | 0.043 | 0.979 | 0.259 | 0.611 | 1.600 | 0.449 | 0.414 | 0.520 | 0.080 | 0.499 |
| SRL | 0.022 | 0.883 | 0.344 | 0.842 | 0.325 | 0.569 | 0.697 | 0.706 | 0.284 | 0.594 | 0.012 | 0.498 |
| SM | 8.499 | **0.004** | 0.546 | 0.761 | 0.471 | 0.493 | 21.479 | **<0.001** | 2.641 | 0.104 | 0.256 | 0.555 |
| $\delta^{13}C$ | 19.695 | **<0.001** | 0.331 | 0.847 | 0.479 | 0.489 | 0.893 | 0.640 | 2.677 | 0.102 | 0.193 | 0.489 |
| FFD | 8.011 | **0.005** | 0.064 | 0.968 | 0.658 | 0.417 | 3.244 | 0.198 | 0.002 | 0.962 | 0.084 | 0.495 |
| LI | 11.838 | **0.001** | 0.056 | 0.972 | 0.534 | 0.465 | 3.385 | 0.184 | 0.579 | 0.447 | 0.092 | 0.555 |
| PC1 | 0.037 | 0.848 | 0.385 | 0.825 | 0.348 | 0.555 | 7.797 | **0.020** | 0.051 | 0.821 | 0.047 | 0.545 |
| PC2 | 13.509 | **<0.001** | 0.121 | 0.941 | 0.362 | 0.547 | 5.817 | 0.055 | 0.800 | 0.371 | 0.153 | 0.494 |
| PC3 | 4.920 | **0.027** | 0.085 | 0.959 | 0.184 | 0.668 | 7.970 | **0.019** | 0.453 | 0.501 | 0.089 | 0.594 |

**Table S6.** Effects of interspecific trait difference (TD, $|Trait_{focal} – Trait_{competitor}|$), site (low, middle or high), the origin of competitors (current or novel) and their interactions on niche overlap (*n* = 93 across all species and sites). The effects were tested using mixed-effects models for each trait, with species as random effects. P values smaller than 0.05 are in boldface. See Table 1 for trait abbreviations.

| **Trait** | **TD** | | **Site** | | **Origin** | | **TD:Site** | | **TD:Origin** | | $R^2_{marg}$ | $R^2_{cond}$ |
|---|---|---|---|---|---|---|---|---|---|---|---|---|
| | *d.f.* = 1 | | 2 | | 1 | | 2 | | 1 | | | |
| | *F* | *P* | *F* | *P* | *F* | *P* | *F* | *P* | *F* | *P* | | |
| RH | 1.114 | 0.291 | 2.239 | 0.326 | 5.594 | **0.018** | 0.788 | 0.674 | 0.008 | 0.930 | 0.033 | 0.730 |
| VH | 0.436 | 0.509 | 2.048 | 0.359 | 4.407 | **0.036** | 0.459 | 0.795 | 0.247 | 0.619 | 0.032 | 0.722 |
| RMF | 0.785 | 0.376 | 2.575 | 0.276 | 5.090 | **0.024** | 0.976 | 0.614 | 1.059 | 0.304 | 0.038 | 0.730 |
| LA | 8.779 | **0.003** | 3.296 | 0.192 | 8.293 | **0.004** | 2.496 | 0.287 | 1.918 | 0.166 | 0.112 | 0.770 |
| LDMC | 18.861 | **<0.001** | 4.049 | 0.132 | 2.874 | 0.090 | 0.075 | 0.963 | 1.072 | 0.300 | 0.107 | 0.778 |
| SLA | 0.442 | 0.506 | 1.984 | 0.371 | 3.680 | 0.055 | 0.313 | 0.855 | 0.027 | 0.870 | 0.023 | 0.752 |
| LN | 2.861 | 0.091 | 2.169 | 0.338 | 3.483 | 0.062 | 5.389 | 0.068 | 0.000 | 0.990 | 0.060 | 0.724 |
| LC | 0.090 | 0.764 | 2.129 | 0.345 | 4.821 | **0.028** | 0.149 | 0.928 | 1.135 | 0.287 | 0.027 | 0.737 |
| FRP | 0.795 | 0.373 | 2.444 | 0.295 | 4.722 | **0.030** | 0.764 | 0.683 | 2.415 | 0.120 | 0.070 | 0.762 |
| FRD | 1.767 | 0.184 | 2.130 | 0.345 | 5.291 | **0.021** | 1.625 | 0.444 | 0.000 | 0.990 | 0.047 | 0.730 |
| SRL | 6.296 | **0.012** | 3.881 | 0.144 | 4.624 | **0.032** | 1.091 | 0.579 | 0.320 | 0.572 | 0.066 | 0.793 |
| SM | 4.578 | **0.032** | 1.471 | 0.479 | 2.310 | 0.129 | 1.802 | 0.406 | 0.000 | 0.991 | 0.087 | 0.772 |
| $\delta^{13}C$ | 0.010 | 0.921 | 1.785 | 0.410 | 4.854 | **0.028** | 2.284 | 0.319 | 0.406 | 0.524 | 0.037 | 0.745 |
| FFD | 0.001 | 0.972 | 1.910 | 0.385 | 3.714 | 0.054 | 9.936 | **0.007** | 2.458 | 0.117 | 0.050 | 0.786 |
| LI | 0.337 | 0.562 | 2.115 | 0.347 | 5.591 | **0.018** | 3.246 | 0.197 | 4.061 | **0.044** | 0.049 | 0.766 |
| PC1 | 0.031 | 0.860 | 1.906 | 0.386 | 4.562 | **0.033** | 1.066 | 0.587 | 0.312 | 0.576 | 0.031 | 0.724 |
| PC2 | 5.647 | **0.017** | 3.160 | 0.206 | 3.271 | 0.071 | 0.349 | 0.840 | 0.784 | 0.376 | 0.042 | 0.780 |
| PC3 | 0.001 | 0.979 | 1.981 | 0.371 | 4.685 | **0.030** | 0.408 | 0.815 | 0.289 | 0.591 | 0.026 | 0.732 |

**Figure S1.** Trait values of each species in each site. Each panel is a trait with points and error bars representing the averages and standard deviations of trait values at each site (low, orange; green, middle; blue, high), except for flowering phenology with points and error bars indicating estimated peak, first and last flowering dates (see Methods). Species on the left of the solid line are highland species (*n* = 6), and species on the right are lowland species (*n* = 7). See Table 1 for trait units and see Table S1 for species codes.

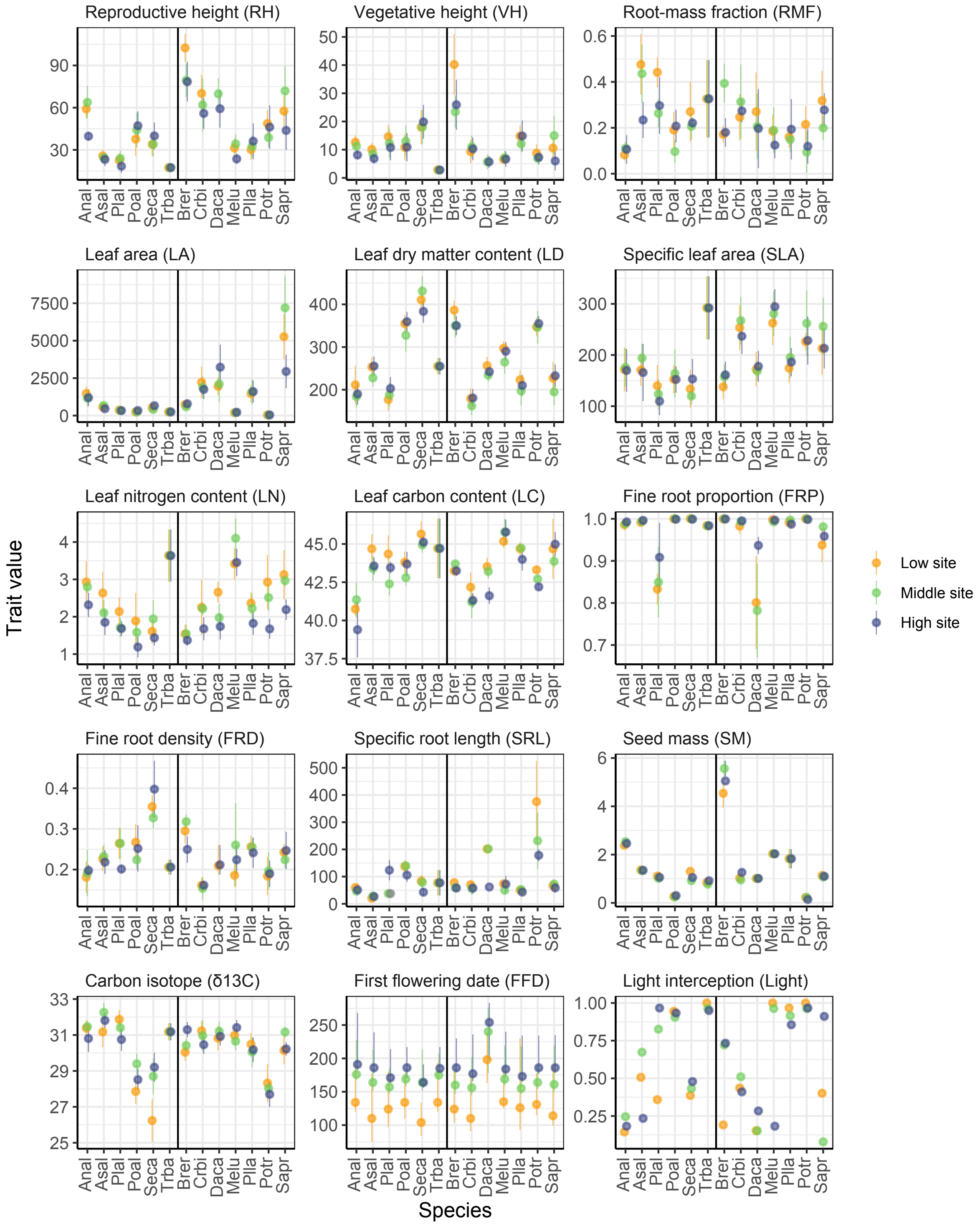

**Figure S2**. Comparison of reproductive height measured on the background plants (i.e. species-level mean) and focal plants (i.e. competitor-specific trait values). Each panel represents a focal species in a site. The horizontal black lines represent the mean reproductive height measured on background plants that were used in the analyses (i.e. species-level mean), while dots with error bars (+/- SD) represent those measured on focal plants.

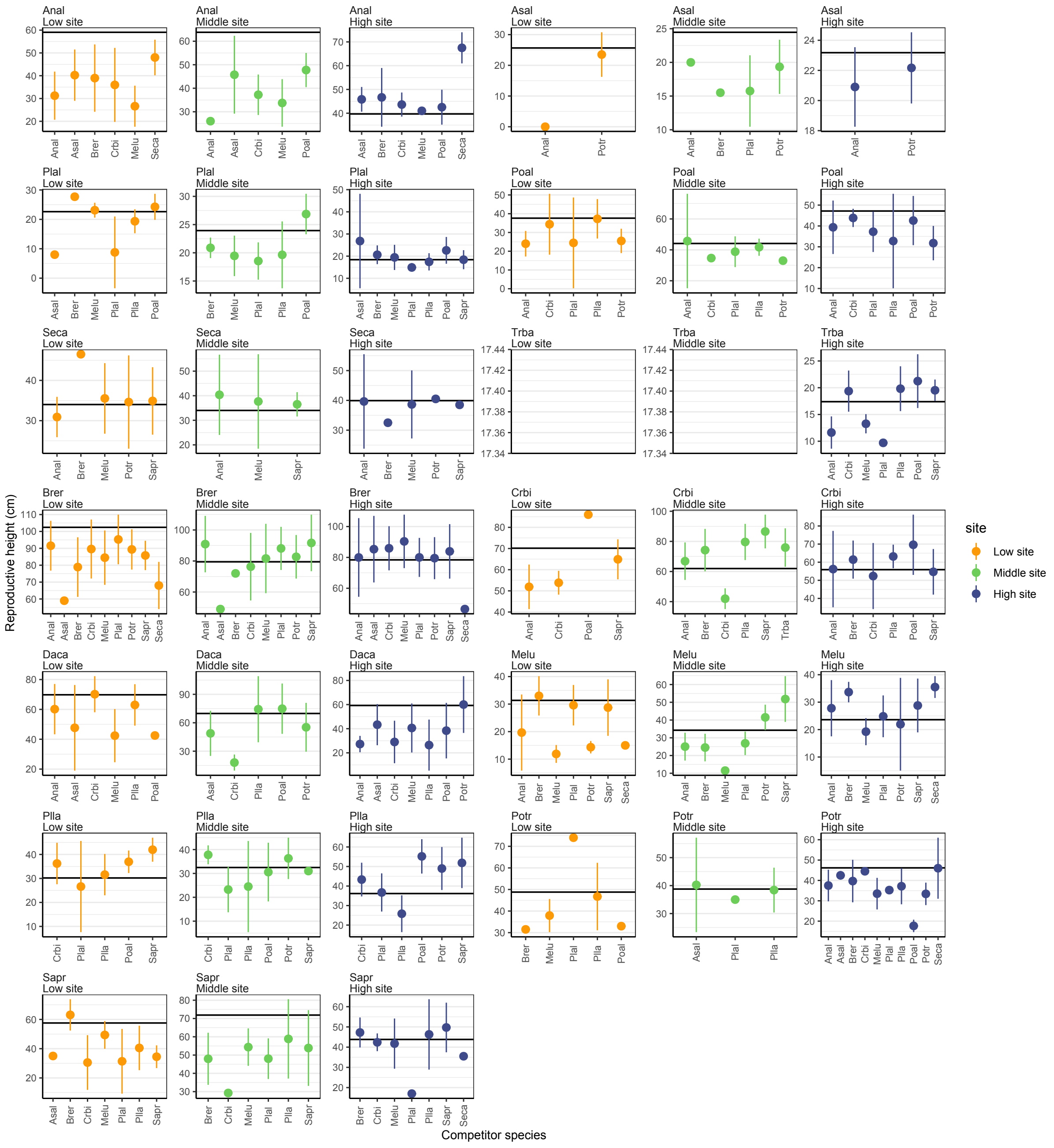

**Figure S3.** Principal components analysis of species' traits. Principal component axes 1 and 2 are shown in panels a and b, and axes 1 and 3 are shown in panels c and d. In panels a and c, arrows indicate the loadings of each trait on principal component axes (see Table 1 for trait abbreviations). Panels b and d show the distribution of species based on their PCA axis values in the three sites (low site, orange; middle site, green; high site, blue) for highland (circled) and lowland species.

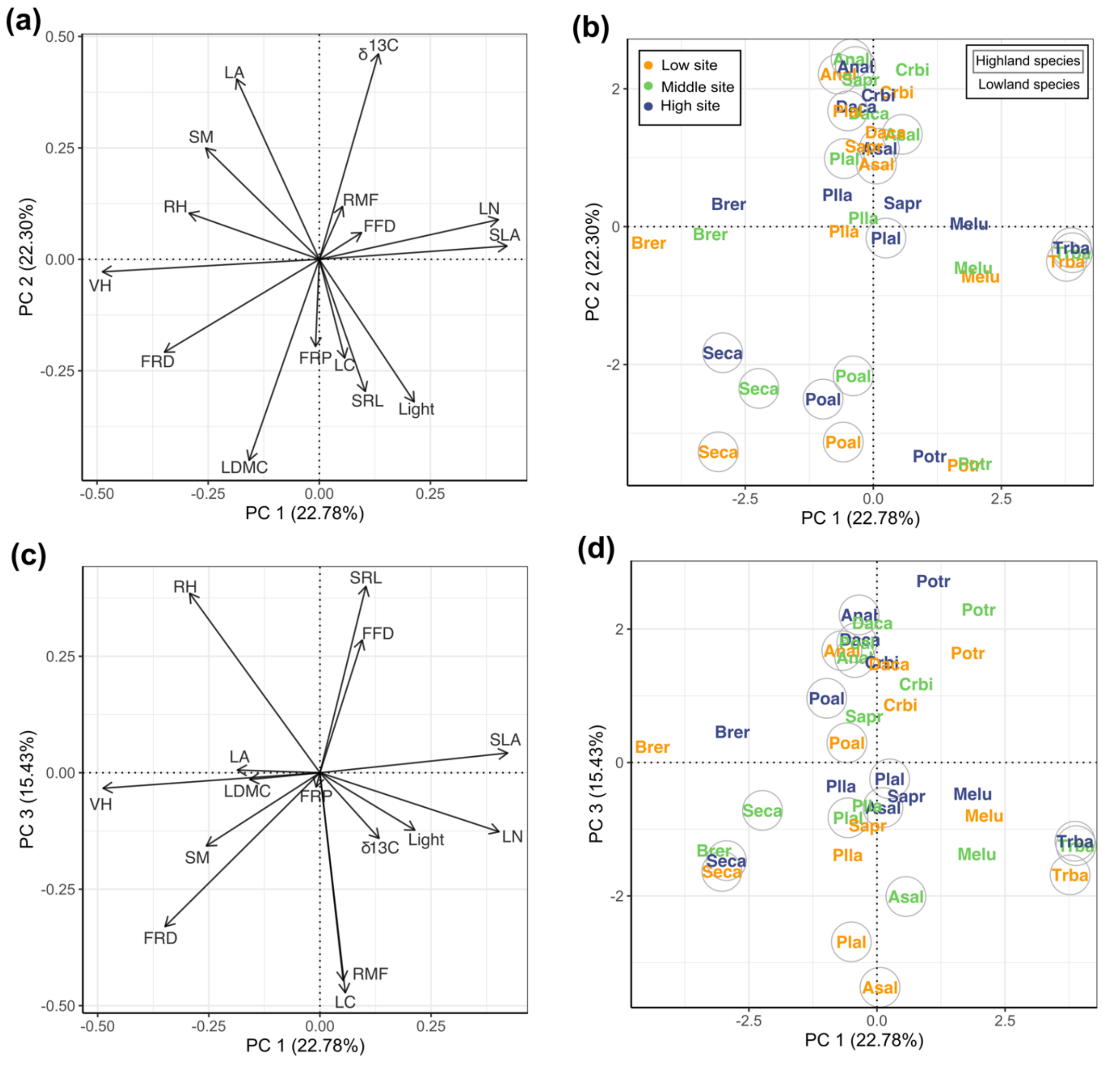

**Figure S4**. The interspecific trait differences (i.e. $|\text{Trait}_{focal} – \text{Trait}_{competitor}|$) among current and novel species pairs in the three sites across an elevation gradient (low, 890 m; middle, 1400 m; high, 1900 m). Each panel represents one trait, boxplots represent the median, first, and third quartiles, respectively, summarized across species pairs within each site, while the upper and lower whiskers indicate 1.5 times the first and third quartiles, respectively. See Table 1 for trait abbreviations and Table S3 for statistical tests.

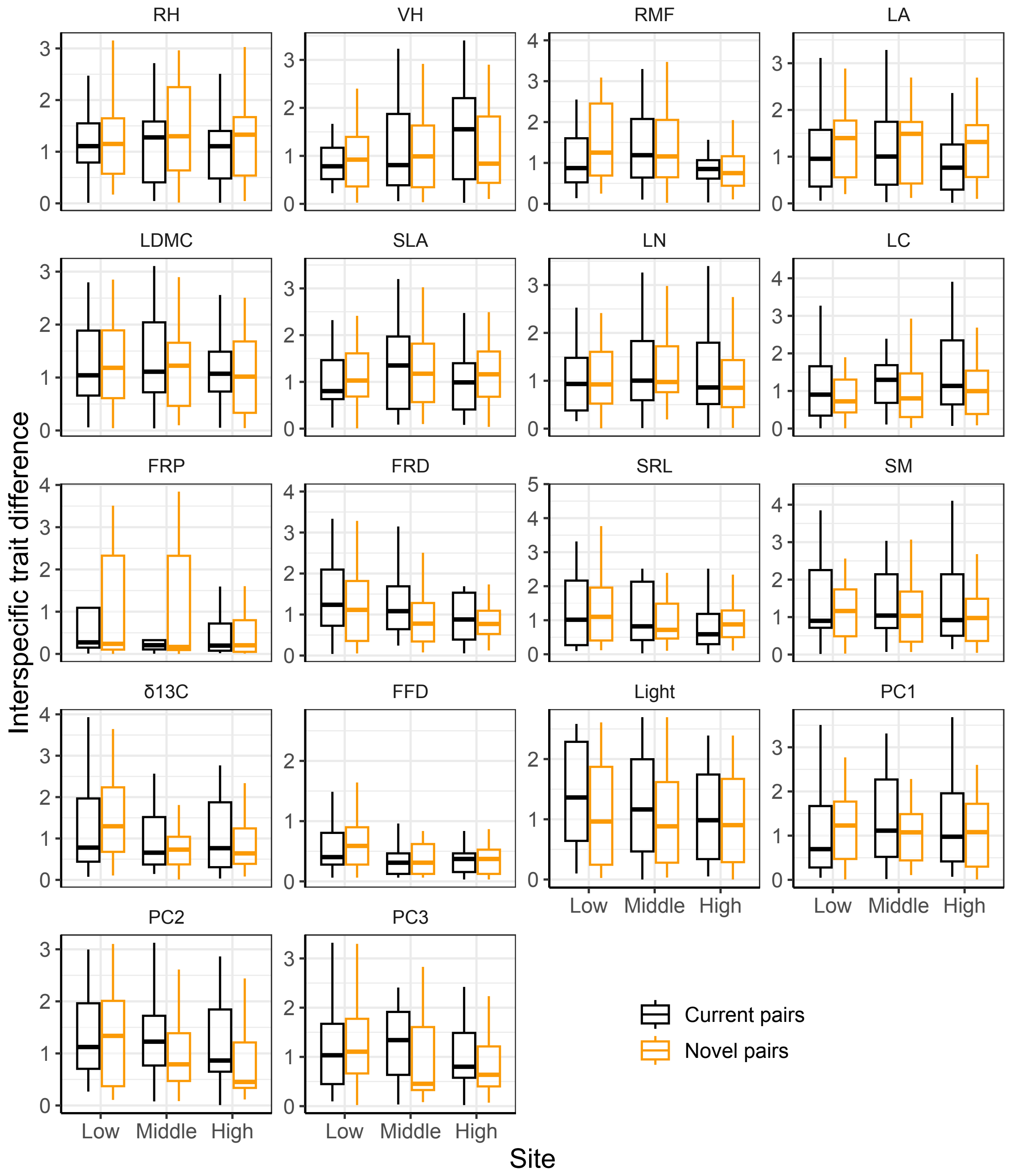

**Figure S5**. The intensity of interactions as measured as response ratio (i.e., $\lambda_{invasion} / \lambda_{intrinsic}$, on a log scale) for current (black) and novel pairs (grey) in the three sites across an elevation (low, 890 m; middle, 1400 m; high, 1900 m). Positive values of log response ratios indicate facilitation, while negative values indicate competition. Boxplots represent the median, first, and third quartiles summarized across species pairs within each site, while the upper and lower whiskers indicate 1.5 times the first and third quartiles, respectively. The strength of species interactions was similar across the elevation gradient ($F_{2, 250}$ = 0.653, P = 0.721) and tended to be greater (i.e. more negative) between novel than current pairs ($F_{1, 250}$ = 3.305, P = 0.069), based on a mixed-effects model with site (low, middle or high) and origin of competitors (current or novel) and their interaction as fixed effects and the identity of focal (*n* = 13) and competitor (*n* = 12) species as random effects. After excluding facilitation (positive log-transformed response ratios; *n* = 20), competition intensity was higher for novel than current pairs ($F_{1, 230}$ = 4.195, P = 0.041) but still similar across the sites ($F_{2, 230}$ = 3.918, P = 0.141).

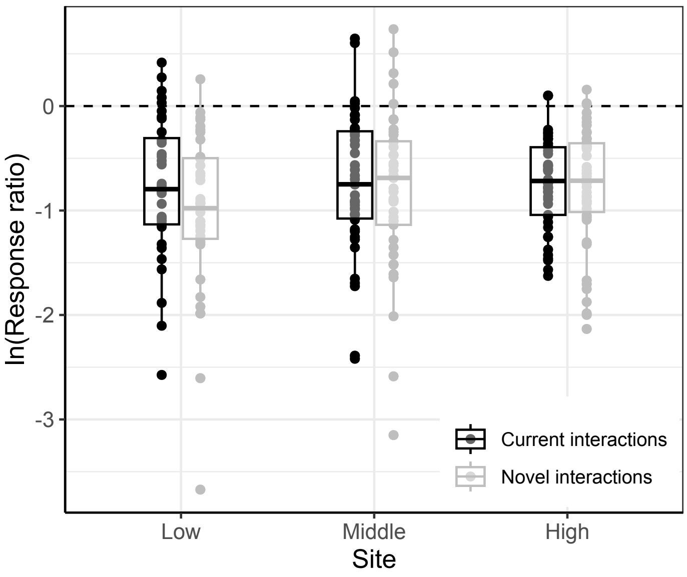

**Figure S6**. The invasion population growth rate (on a logarithm scale) for current (black) and novel pairs (grey) in the three sites across an elevation (low, 890 m; middle, 1400 m; high, 1900 m). Boxplots represent the median, first, and third quartiles summarized across species pairs within each site, while the upper and lower whiskers indicate 1.5 times the first and third quartiles, respectively. Invasion growth rate of the focal species was similar across the elevation gradient ($F_{2,\ 250}$ = 3.509, P = 0.173) and when they interacted with current and novel competitors ($F_{1,\ 250}$ = 2.353, P = 0.125), based on a mixed-effects model with site (low, middle or high) and origin of competitors (current or novel) and their interaction as fixed effects and the identity of focal (*n* = 13) and competitor (*n* = 12) species as random effects. Notably, the variation in invasion growth rates appeared to decrease towards higher elevations.

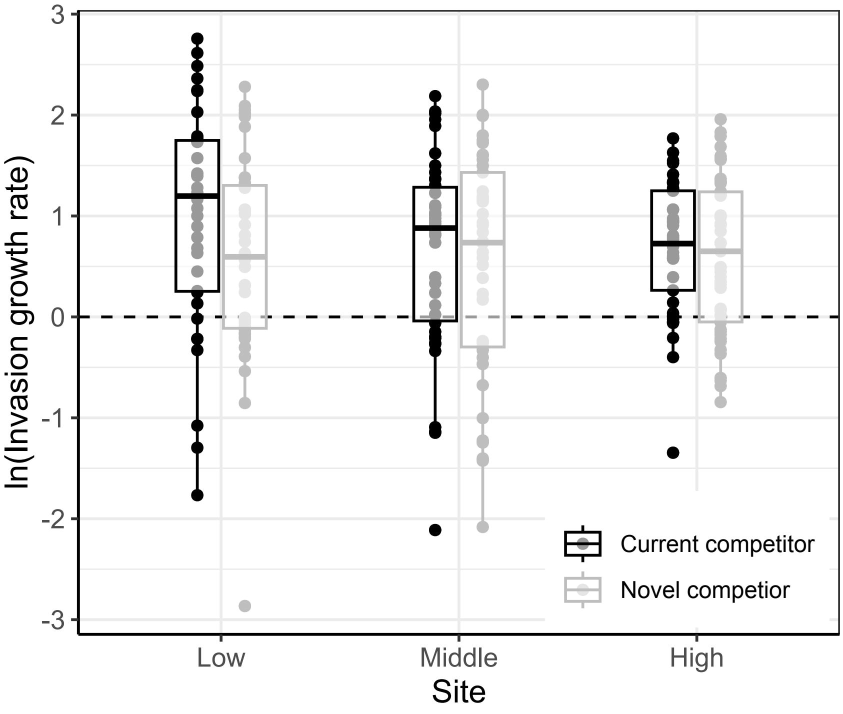

**Figure S7**. Relationships between invasion growth rates (on a log scale) and interspecific trait differences (i.e., $trait_{focal} – trait_{competitor}$) in the three study sites across an elevation gradient (low, orange; green, middle; blue, high). Each panel represents a trait. Each point represents a species pair. Lines represent fitted relationships, with solid lines indicating significant relationships (95% CIs of the slopes did not include zero) and dashed lines indicating non-significant relationships. The significance of trait hierarchy x site interactions is also shown for each trait (* P < 0.05; n.s. P > 0.05; Table 2).

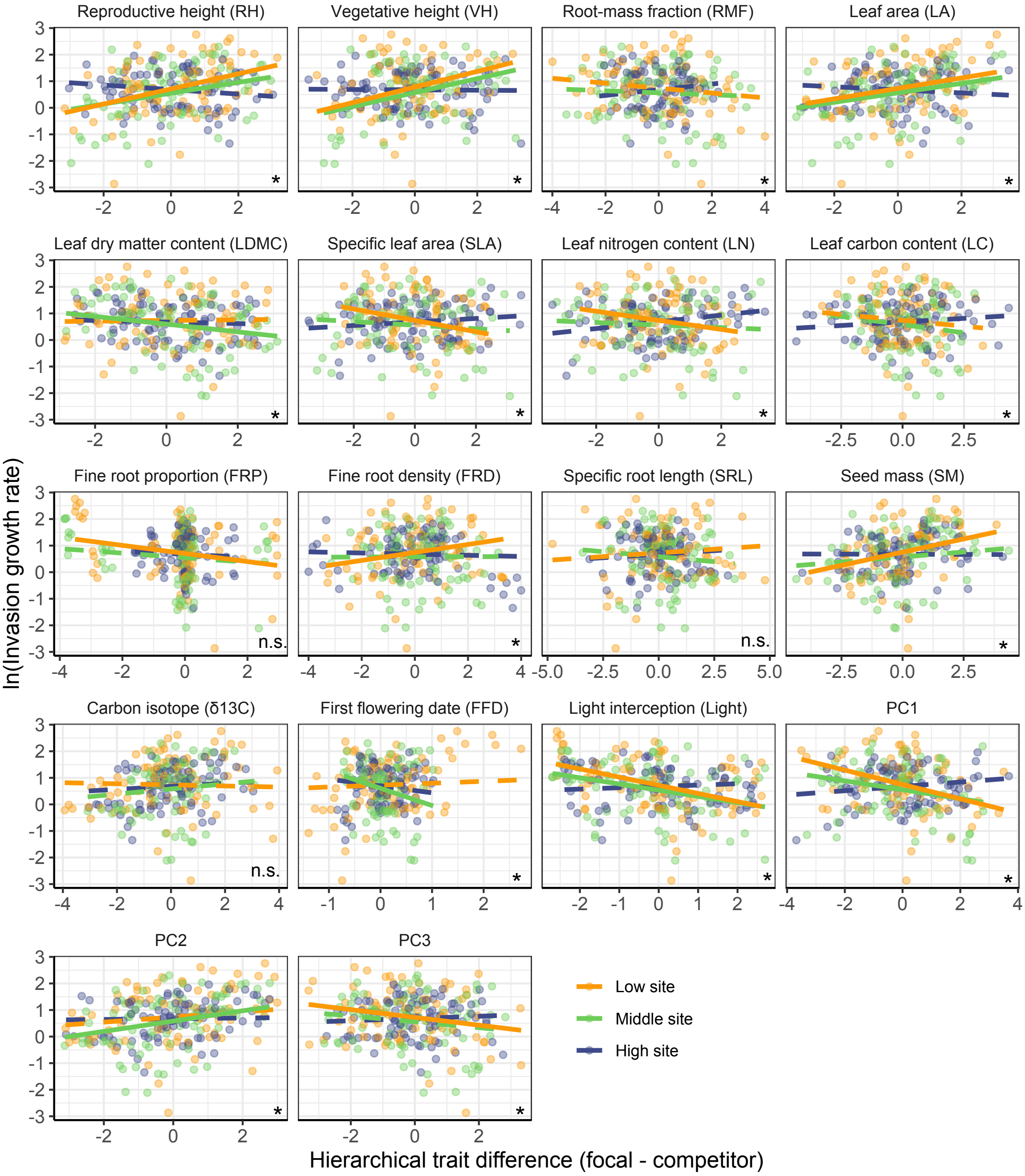

**Figure S8**. Effect sizes of trait differences on invasion growth rates (a, d), relative fitness differences (b, e) and niche overlap (c, f) for current (the first row, black) and novel (the second row, orange) interactions across the elevation gradient. Effect sizes are indicated with dots, and the lines connecting the three effect size values of each trait across the three study sites show the trend of variation along the elevation gradient.

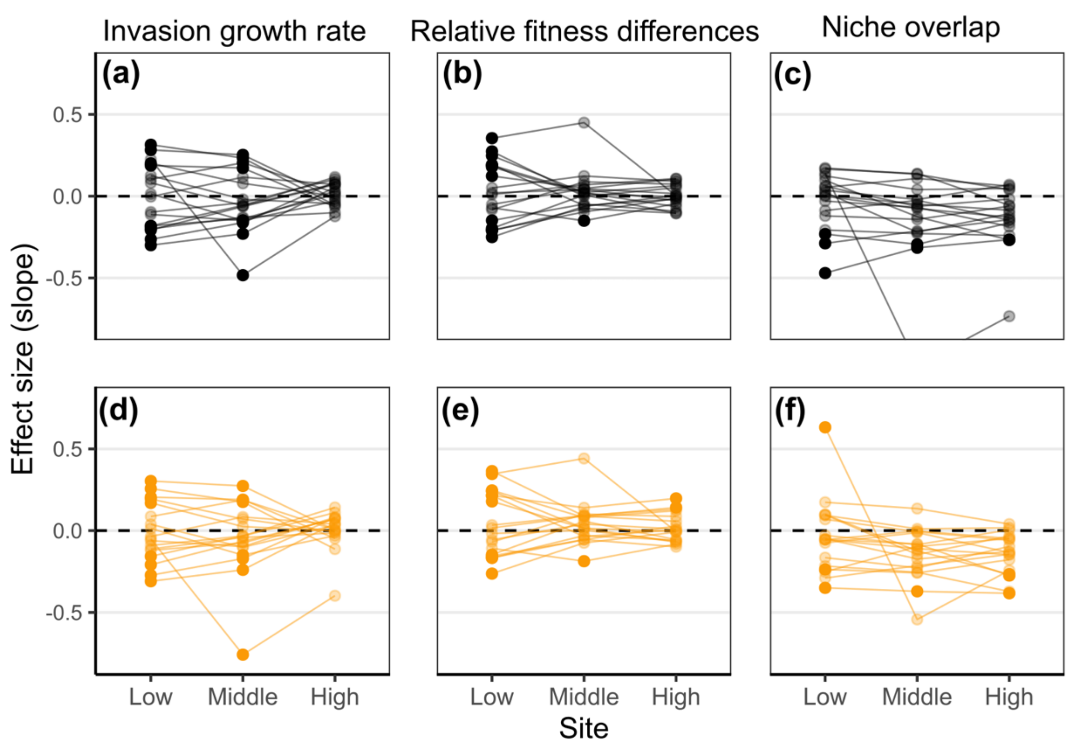

**Figure S9**. Relationships between the number of traits used to calculate trait differences (i.e. $Trait_{focal} – Trait_{competitor}$) and deltaAICc (a) and effect size (b) across the three sites. deltaAICc and slopes are obtained from the models with invasion growth rate (IGR) on a logarithm scale as response variables and multiple trait differences as explanatory variables. The multiple trait differences were measured as the Euclidean distance in the multiple trait space. The models were fitted separately at each site (n = 75, 83 and 92 at the low, middle and high sites, respectively).

**(a)**

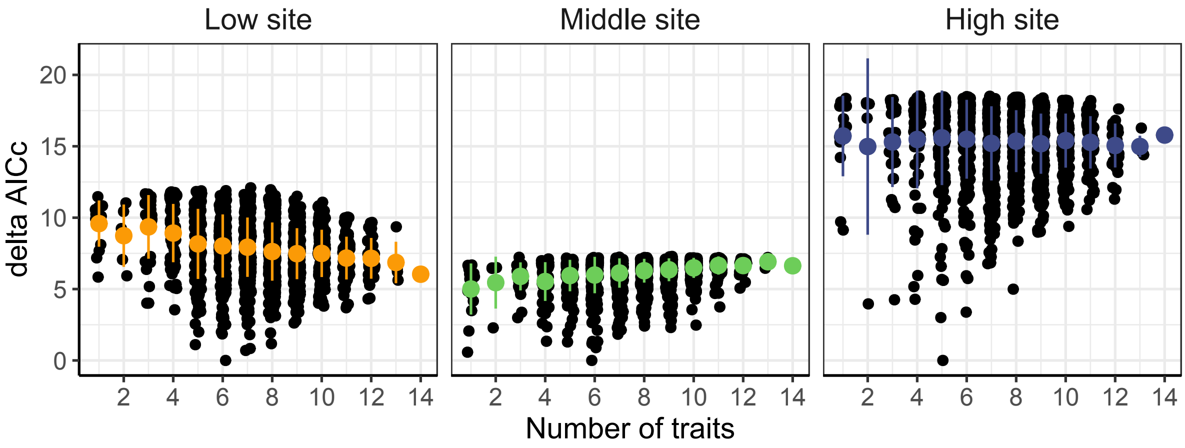


**(b)**

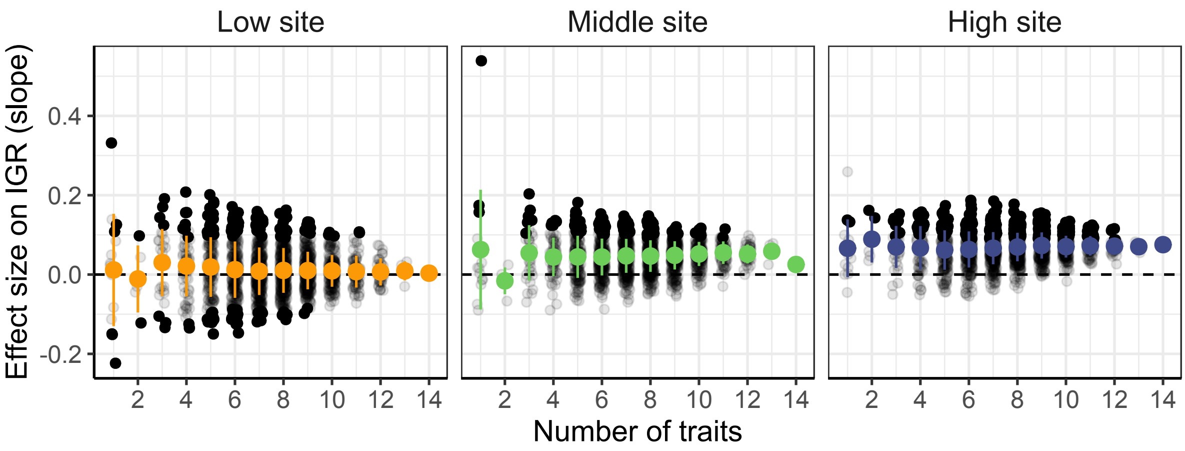

**Figure S10**. Relationships between the number of traits used to calculate trait differences (i.e. $Trait_{focal} – Trait_{competitor}$) and deltaAICc (a) and effect size (b) across the three sites. In contrast to Fig. S9, deltaAICc and slopes are obtained from the models with relative fitness differences (RFD) on a logarithm scale as response variables and multiple trait differences as explanatory variables. The multiple trait differences were measured as the Euclidean distance in the multiple trait space. The models were fitted separately at each site (n = 29, 28 and 36 at the low, middle and high sites, respectively).

**(a)**

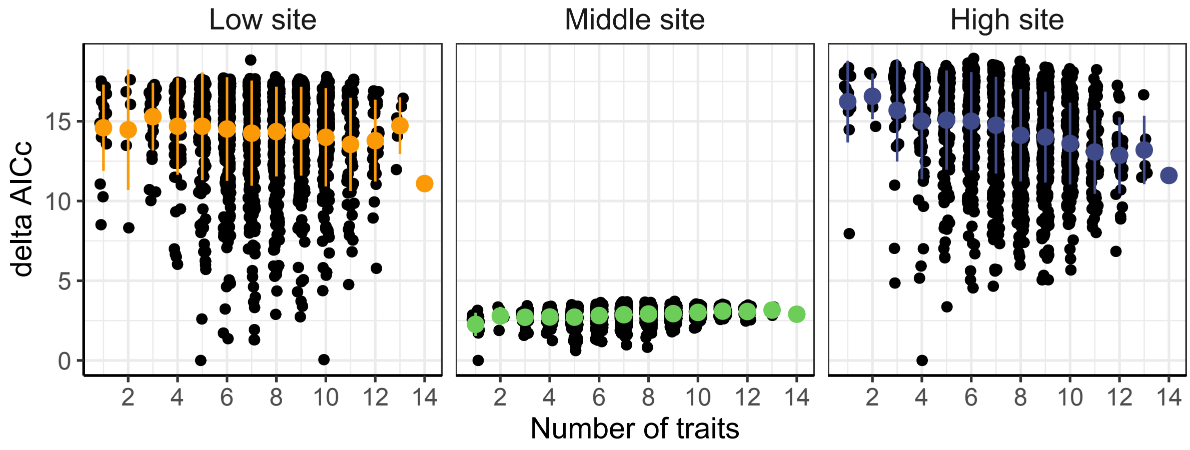


**(b)**

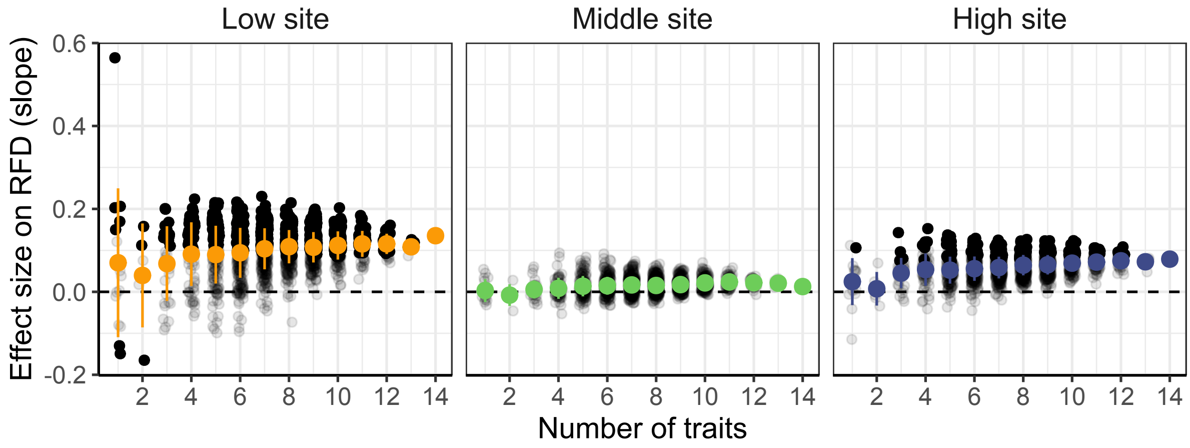

**Figure S11**. Relationships between the number of traits used to calculate trait absolute differences (i.e. |$\text{Trait}_{\text{focal}}$ – $\text{Trait}_{\text{competitor}}$|) and deltaAICc (a) and effect size (b) across the three sites. In contrast to Fig. S9, deltaAICc and slopes are obtained from the models with niche overlap (i.e., 1- niche difference) on a logarithm scale as response variables and multiple trait differences as explanatory variables. The multiple trait differences were measured as the Euclidean distance in the multiple trait space. The models were fitted separately at each site (n = 29, 28 and 36 at the low, middle and high sites, respectively).

**(a)**

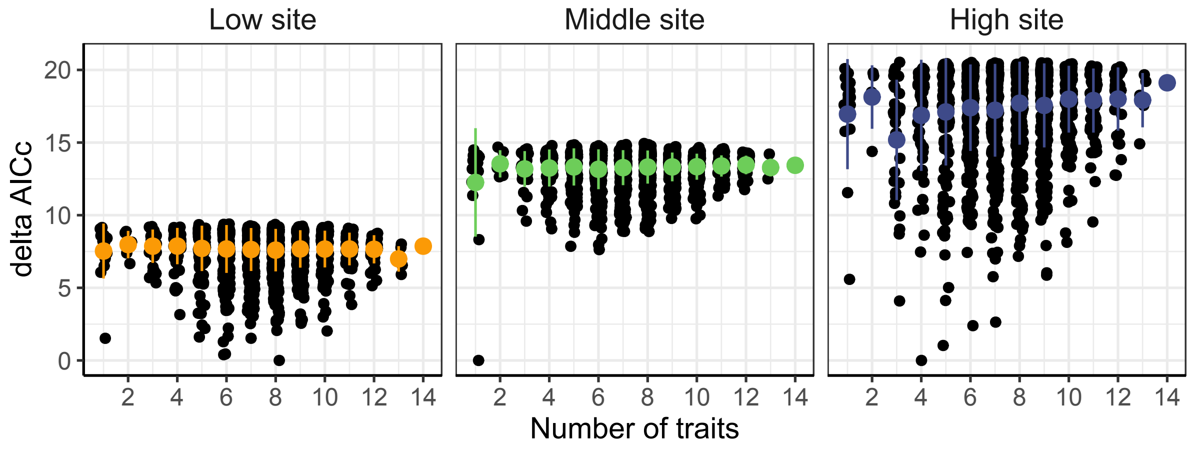


**(b)**

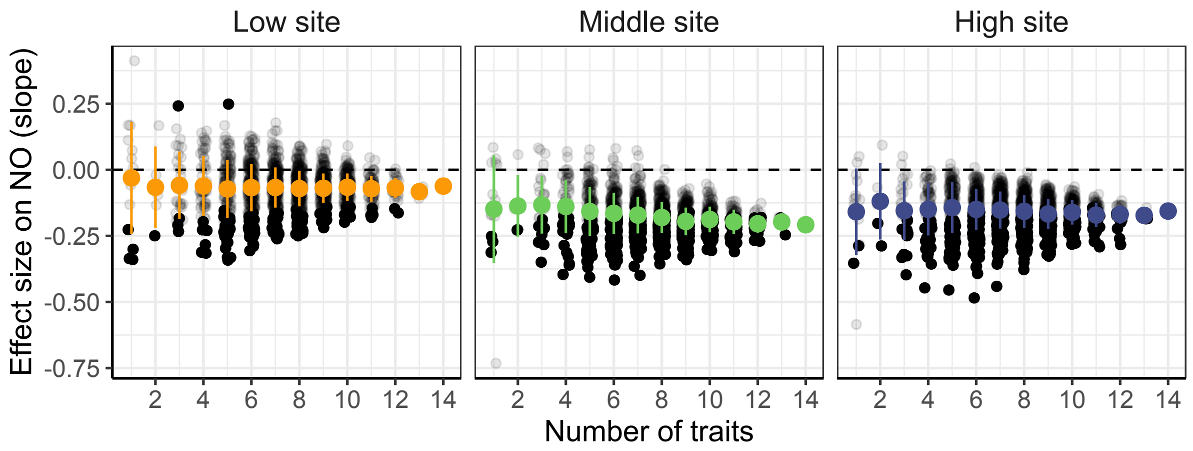

**Figure S12**. Relationships between interspecific differences in reproductive height (i.e. TD = $\text{Height}_{focal}$ – $\text{Height}_{competitor}$) and invasion growth rate on a logarithm scale. In the panel a, TD was calculated using the speciesl mean same as the in the main text. In the panel b, TD was calculated using the competitor-specific height of focal plants (Fig. S2).

**(a)**

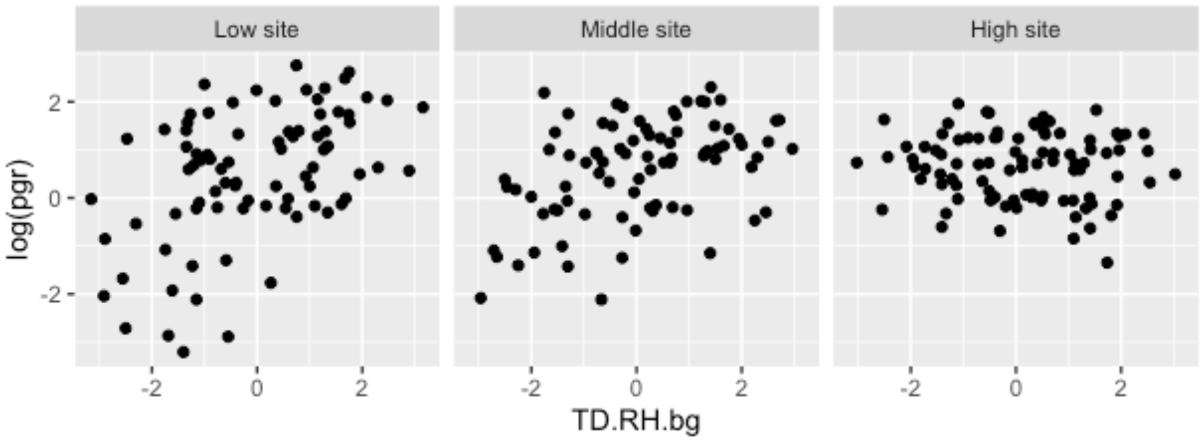


**(b)**

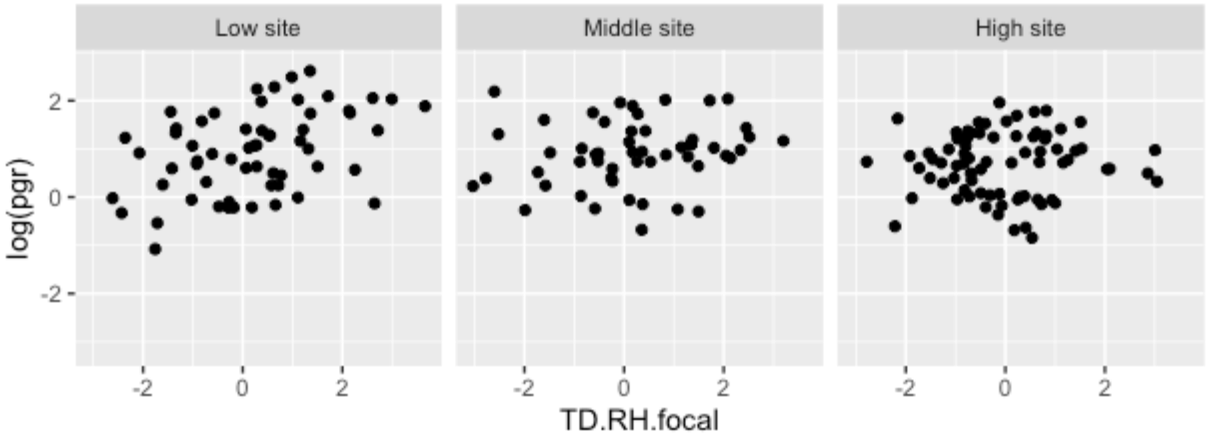
